# Combined small molecule treatment accelerates timing of maturation in human pluripotent stem cell-derived neurons

**DOI:** 10.1101/2022.06.02.494616

**Authors:** Emiliano Hergenreder, Yana Zorina, Zeping Zhao, Hermany Munguba, Elizabeth L. Calder, Arianna Baggiolini, Andrew P. Minotti, Ryan M. Walsh, Conor Liston, Joshua Levitz, Ralph Garippa, Shuibing Chen, Gabriele Ciceri, Lorenz Studer

## Abstract

The maturation of human pluripotent stem cell (hPSC)-derived neurons mimics the protracted timing of human brain development, extending over months and years to reach adult-like function. Prolonged *in vitro* maturation presents a major challenge to stem cell-based applications in modeling and treating neurological disease. We designed a high-content imaging assay based on morphological and functional readouts in hPSC-derived cortical neurons to reveal underlying pathways and to identify chemicals capable of accelerating neuronal maturation. Probing a library of 2688 bioactive drugs, we identified multiple compounds that drive neuronal maturation including inhibitors of LSD1 and DOT1L and activators of calcium-dependent transcription. A cocktail of 4 factors **G**SK-2879552, **E**PZ-5676, **N**MDA and Bay **K** 8644, which we collectively termed GENtoniK, triggered maturation across all assays tested including measures of synaptic density, electrophysiology and transcriptomics. Remarkably, GENtoniK was similarly effective in enhancing neuronal maturation in 3D cortical organoids and in spinal motoneurons, and improved aspects of cell maturation in non-neural lineages such as melanocytes and pancreatic beta cells. These results demonstrate that the maturation of multiple hPSC-derived cell types can be enhanced by simple pharmacological intervention and suggests that some of the mechanisms controlling the timing of human maturation are shared across lineages.

## Introduction

Recent advances in human pluripotent stem cell (hPSC) differentiation enable the derivation of a myriad of specific subtypes of neurons on demand^1–3^. However, the application of this technology remains hampered by the slow maturation rates of human cells resulting in prolonged culture periods for the emergence of disease-relevant phenotypes. Indeed, most neurological and psychiatric disorders manifest as impairments in postnatal or adult neuron functions such as synaptic connectivity^4^, dendritic arborization^5^, and electrophysiological function^6^. Therefore, developing strategies to accelerate the maturation of hPSC-derived neurons is critical to realize their full potential in modeling and treating neural diseases.

Multiple cell-extrinsic factors have been identified as contributors to neuron maturation, including glial cells^7^, network activity^8^ and neurotrophic factors^9^. However, within a given micro-environment, cell-intrinsic maturation rates appear dominant and seem to be determined by a species-specific molecular clock, which runs especially slow in human neurons^10,11^. For example, the maturation of hPSC-derived cortical neurons transplanted into the developing mouse brain follows human-specific timing, requiring 9 months to achieve mature, adult-like morphologies and spine function^12^. Similarly, the transplantation of mouse, versus pig versus human midbrain dopamine neurons into the brain of Parkinsonian rats results in graft-induced functional rescue after 4 weeks, 3 months or 5 months respectively, indicating that transplanted cells retain their intrinsic, species-specific *in vivo* maturation timing rather than adopting the timing of the host speies^13^.

Here we aimed at identifying effectors of neuronal maturation and developing a chemical strategy to accelerate it. We designed a multi-phenotypic, image-based assay to monitor maturation in nearly pure populations of hPSC-derived deep layer cortical neuron cultures and applied it to screen 2688 bioactive compounds. Among the screening hits, compounds targeting chromatin remodeling and calcium-dependent transcription were combined into a maturation cocktail that was effective across a broad range of maturation phenotypes and capable of driving aspects of maturation in both neuronal and non-neuronal lineages.

## Results

### High content assay of neuron maturity

The phenotypic complexity of neurons makes single-readout assays unsuitable to fully capture maturation stages. Therefore, we used a multi-phenotype approach (via high-content screening, HCS)^14^ to design an assay that simultaneously monitors distinct features of neuronal maturation (Fig. 1a). Dendritic outgrowth is a widely used parameter of neuron maturity^15^ and can be monitored through automated tracing of microtubule-associated protein 2 (MAP2) immunostaining (Fig. 1b, c). Changes in nuclear size and morphology are also characteristic of neuron development and maturation^16^ and can be tracked via DAPI counterstaining (Fig. 1b, c). As an indirect measurement of neuronal function and excitability, we quantified the nuclear expression of immediate early gene (IEG) products FOS and EGR1 following 2 hours of KCl stimulation (Fig. 1b, d). IEGs are defined by their rapid induction in the absence of de-novo protein synthesis by stimuli that include sustained membrane depolarization in neurons^17^. In contrast to more traditional measures of neuronal activity such as calcium imaging and electrophysiology, IEG immunoreactivity is readily scalable as a readout for thousands of treatment conditions. However, IEGs can be triggered by stimuli other than neuronal activity including growth factor signaling^18^ and cellular stress responses^19^. Therefore, to avoid direct activation of IEGs, we used transient compound treatment (day 7-14) and performed all measurements after rinsing of compounds followed by culture in compound-free medium for an additional 7 days (day 14-21) prior to analysis (Fig. 1a). Furthermore, we recorded IEGs under both basal and KCl-stimulated conditions to specifically determine the depolarization-induced signal by subtracting baseline from KCl-induced responses. Measuring maturation readouts only after compound withdrawal enabled the identification of compounds that trigger a long-lasting “memory” of a maturation stimulus even after compound removal.

**Fig. 1.**
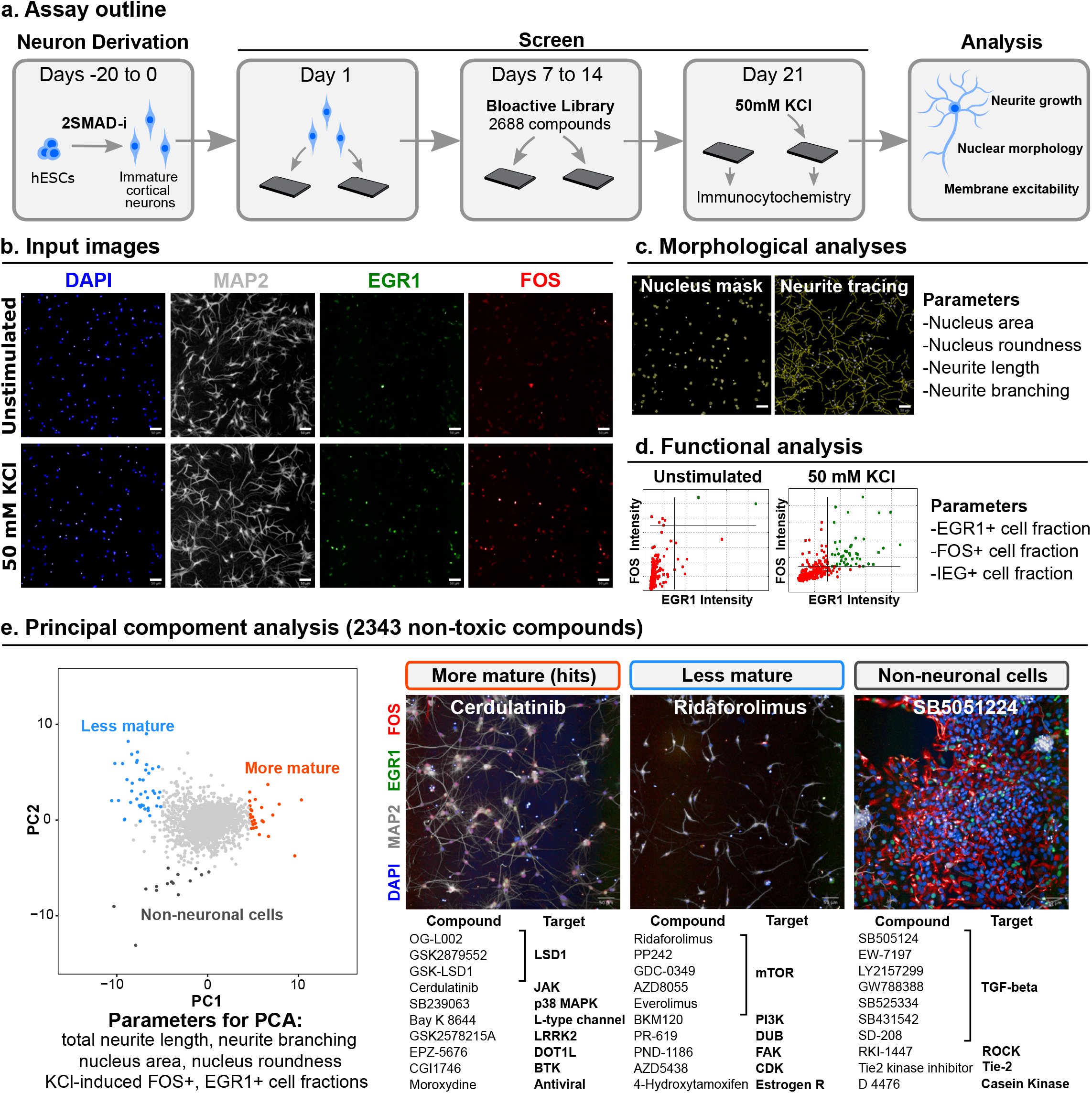
High-content chemical screen for drivers of neuron maturation. **a**, Outline of screening protocol in hPSC-derived excitatory cortical neurons. 2SMAD-i, dual SMAD inhibition. **b**, Example of input immunofluorescent images. Top: unstimulated neurons at day 21 post plating. Bottom, neurons received 50 mM of KCl 2 hours before fixation. **c**, automated analysis of neuron morphology. Left, nuclei detection mask from DAPI channel. Right, automated neurite tracing from MAP2 channel. **d**, Quantification of neuron excitability by applying an intensity threshold to FOS and EGR1 channels within the nuclear mask. **e**, Principal component analysis of screened compound library computed from 6 maturity parameters (z-scores averaged from n = 2 independent screen runs). Left, PCA plot of 2343 non-toxic library compounds (out of 2688 total compounds tested) with phenotypic clustering of maturation enhancing (orange), maturation inhibiting (blue), and non-neuronal proliferation enhancing (grey) compounds. Right, representative screen images and 10 representative hit compounds within each cluster. Scale bars are 50 μm.

While these readouts are pan-neuronal, and therefore appropriate across different neuronal lineages, we chose cortical neurons for the screen for both technical and biological reasons. Cortical neurons can be derived at high efficiency in the absence of expensive recombinant proteins, and their even cell distribution free of clusters makes them amenable to high-throughput imaging. They also represent a brain region that undergoes a particularly protracted development, and a region of great importance to human neurological disease. Our cortical neuron differentiation protocol yields highly pure populations of post-mitotic deep-layer TBR1+ cells, which can be readily scaled, cryopreserved and directly thawed for use in large-scale assays (Supp. Fig. 1a-d).

To benchmark the assay performance in mature cells, we employed primary embryonic rat cortical neurons, which quickly and reliably develop mature-like functionality *in vitro*^20^. At 14 days after plating, rat neurons displayed large and round nuclei (130 μm^2^, 0.93 roundness index), extensive neurite growth (>2500 μm/ neuron), and near 100% of the neurons showed KCl-induced IEG responses (Supp. Fig. 1e-i). In contrast, in human PSC-derived cortical neurons, these properties only very gradually increased over a 50-day culture period and never reached the maturity of their rodent counterparts (Supp. Fig. 1j-m). These results indicate that our multi-phenotypic assay reliably captures the maturation of developing rat and human PSC-derived human cortical neurons.

### Chemical screen for maturation enhancers

We next applied our maturity assay to screen a library of 2688 bioactive compounds in hPSC-derived cortical neurons (Supp. Fig. 2a). The library was applied at 5 μM and standard scores (z-scores) of duplicate screen runs were averaged for analysis. Viability was determined by quantifying intact nuclei, and 325 toxic compounds with a z-score below −2 were excluded from further analysis (Supp. Fig. 2b). For HCS hit selection, we applied principal component analysis (PCA) to 6 maturity z-scores to identify patterns of distribution among compounds, avoiding single threshold hit discrimination^21^ (Fig. 1e, left panel). The 6 parameters were: nucleus size and roundness, total neurite length and branching (number of segments per cell), and fractions of KCl-induced FOS+ and EGR1+ cells. We identified 3 phenotypic clusters of compounds by PCA: maturation enhancers (hits); maturation suppressors, consisting mostly of inhibitors of the PI3K/AKT/mTOR axis; and inducers of non-neuronal contaminant proliferation, which were highly enriched in TGF-β signaling inhibitors as well as inhibitors of rho-associated protein kinase (ROCK) and other signaling pathways (Fig. 1e, right panel).

We selected 32 compounds within the mature cluster for validation. While PCA identifies compounds with the greatest overall maturation effect, we reasoned that compounds with strong effects on single parameters could also be of interest. We therefore added the top 5 highest scoring compounds for each, total neurite length and double FOS/EGR1 positive cells, excluding compounds already selected by PCA (Supp. Fig. 3a). Because single-parameter readouts are susceptible to false positives, we excluded drugs with known maturation-independent effects, such as microtubule stabilizers docetaxel and paclitaxel. Interestingly, neurite-only hits included several inhibitors of Aurora kinase, in agreement with recent phenotypic screens targeting this phenotype^22,23^. Using these combined criteria, we selected 42 primary hits (Supp. Table 1).

To validate primary hits, the 42 compounds were applied to the maturity assay in triplicates at the screening concentration (5μM) and ranked by their effect on 4 maturity parameters: nucleus size and roundness, total neurite length, and double KCl-induced FOS/EGR1 cells (Supp. Fig. 3b). The 22 compounds with the highest mean normalized score over DMSO across all parameters underwent additional dose-response studies (Fig. 2a) resulting in the identification of 4 compounds with the most pronounced, dose-dependent effects on the mean maturation score (Fig. 2b).

**Fig. 2.**
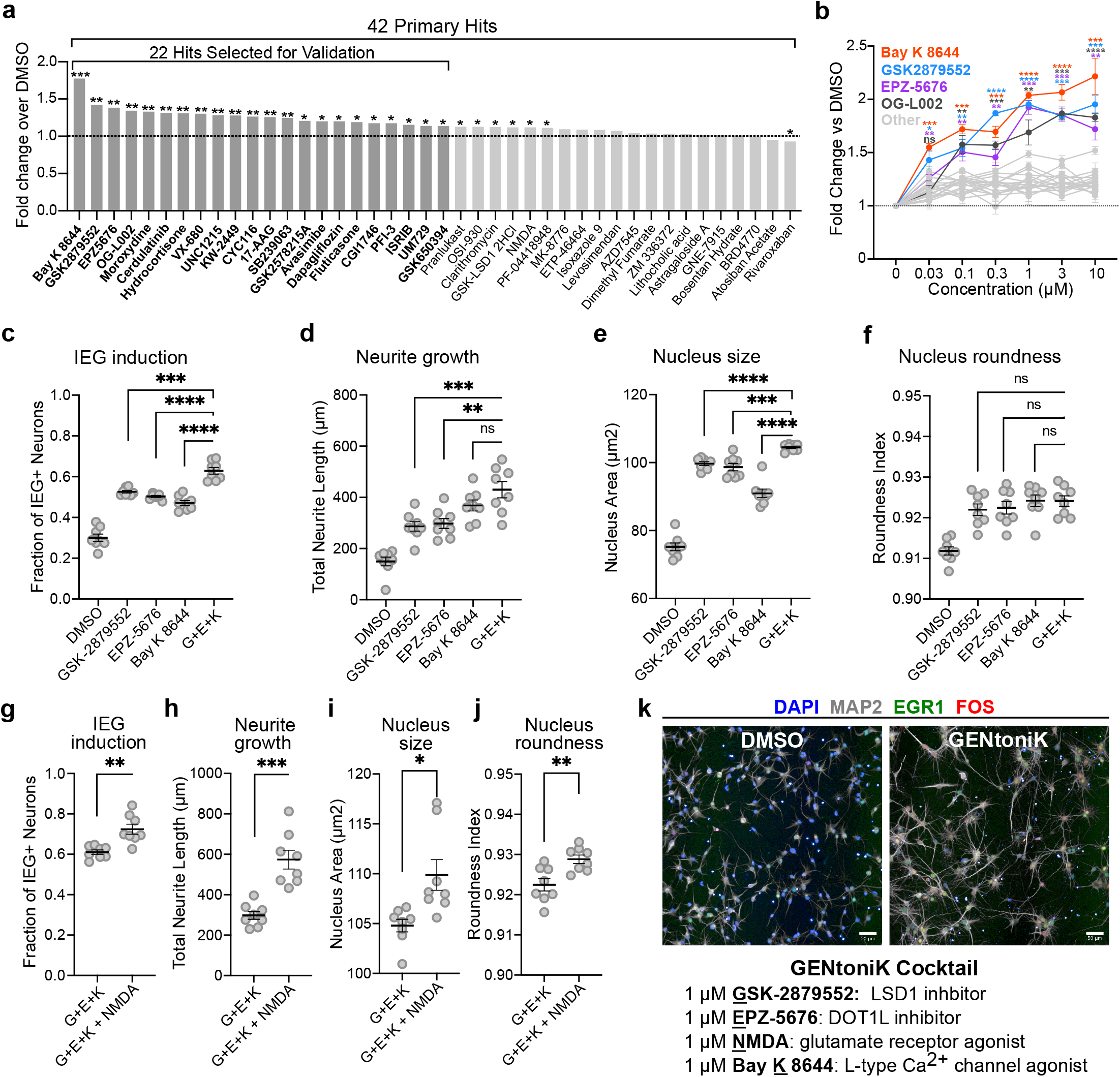
Validation and combination of screen hits identifies maturation-promoting cocktail GENtoniK. **a,** Ranking of primary hits by the mean of 4 maturity parameters (nucleus size and roundness, neurite length, and KCl-induced double FOS/EGR1+ cells) normalized to DMSO (n = 3 microplate wells). 22 top-ranked compounds were selected for validation. **b,** Dose-response validation of 22 screen hits comparing the mean of 4 maturity parameters normalized to DMSO (n = 15 microplate wells from 3 independent differentiation). **c-f**, Comparison of confirmed hits GSK-2879552, EPZ-5676, Bay K 8644, and a combination of the 3 (G+E+K) across maturity parameters IEG induction (**c**), neurite growth (**d**), nucleus size (**e**), and nucleus area (**f**) (n = 8 microplate wells from 2 independent experiments). **g-j**, Comparison of 3-hit drug combination (G+E+K) to the same with the addition of NMDA across maturity parameters IEG induction (**g**), neurite growth (**h**), nucleus size (**i**), and nucleus roundness (**j**) (n = 8 microplate wells from 2 independent differentiations). **k**, Top, representative images of cortical neurons treated with DMSO or maturation promoting cocktail GENtoniK. Bottom, formulation of GENtoniK. **a-b**, Brown-Forsythe and Welch ANOVA with Dunnett’s T3 multiple comparison test. **c-j**, Two-tailed Welch’s *t*-test; asterisks indicate statistical significance. Mean values are represented by a bar graph (**a**) or a line (**c-j**). Error bars represent S.E.M. Scale bars are 50 μm.

### Small molecule cocktail promotes neuron maturity

The 4 confirmed maturation-promoting compounds consisted of two inhibitors of lysine-specific demethylase 1 (LSD1/KDM1A), an inhibitor of disruptor of telomerase-like 1 (DOT1L), and an agonist of L-type calcium channels (LTCC). LSD1 is a histone 3 demethylase at lysine 4 and 9, and a switch of specificity between these 2 substrates has been previously linked to neuron differentiation^24,25^. DOT1L is the sole methyltransferase targeting lysine 79 within the globular domain of histone 3^26^. LTCCs are involved in calcium-dependent transcription and play important roles in neuron development^27^. We reasoned that transcriptional induction by the LTCC agonist might potentiate the effect of chromatin remodeling by epigenetic regulators such as LSD1 and DOT1L. Accordingly, we next sought to determine whether a combination of the hits can further enhance neuron maturation. Because two of the confirmed hits target LSD1, we decided to only pursue one of them (GSK-2879552) for combinatorial experiments, as it displayed a stronger combined effect than OG-L002 (Fig. 2b). A combination of the 3 hit compounds significantly increased IEG induction, neurite growth, and nucleus size, but not nucleus roundness, as compared to the results following single compound treatments (Fig 2c, Supp. Fig. 4a). These effects appear to be independent of cell viability, as neither the individual treatments nor combination significantly altered the number of cells with respect to DMSO (Supp. Fig 4b).

In addition to LTCCs, calcium-dependent transcription is initiated through activation of the NMDA glutamate receptors^28^, which have also been shown to participate in neuron maturation^29^. Interestingly, the compound NMDA itself was among the primary screen hits but did not pass validation as single agent treatment (Fig. 2a). We next tested whether the addition of NMDA could further enhance the maturation parameters in the presence of the above 3 hit combination. We observed significant improvements across all maturity parameters, again without changes in cell survival (Fig. 2d, Supp. Fig. 4c), and we nominated the resulting 4-drug cocktail (**G**SK-2879552, **E**PZ-5676, **N**MDA and Bay **K** 8644) as a maturation-promoting strategy, naming it GENtoniK (Fig. 2e).

### GENtoniK promotes functional neuron maturation

We next validated GENtoniK on additional maturation phenotypes that are orthogonal to those assayed during screening. Establishing independent functional read-outs was particularly important, as three of the proteins targeted by the cocktail have been reported to directly participate in IEG induction in neurons^30–32^. The formation of chemical synapses is a critical step in neuronal development that also occurs in protracted manner in the human cortex^33^. We used immunofluorescent staining in day 35 cortical neurons to assess the effect of GENtoniK on synaptogenesis. Density of synaptic assembly was quantified through the apposition of the pre- and post-synaptic markers SYN1 and PSD95 normalized to dendrite length (Fig. 3a). GENtoniK-treated neurons showed increased density of both pre- and post-synaptic markers per neurite length, as well as an increased density of the apposition of synaptic punctae (Fig. 3b-d).

**Fig. 3.**
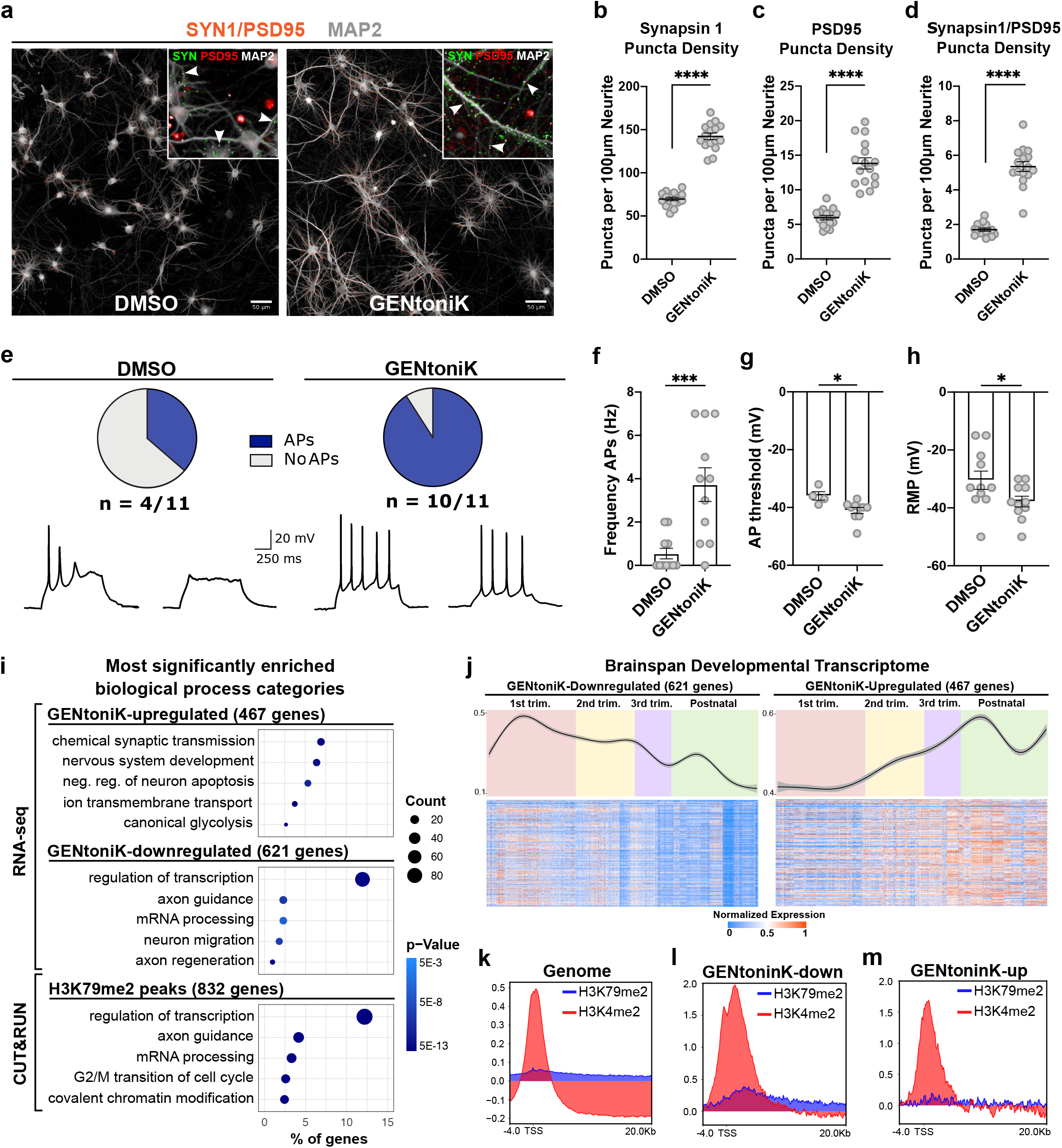
Validation of small molecule maturation strategy with orthogonal readouts. **a**, Representative images for synaptic marker detection in day 35 hPSC-derived cortical neurons that received DMSO versus GENtoniK treatment from days 7 to 21. Orange dots represent instances of SYN1 and PSD95 apposition. Inset, input immunofluorescent images used for quantification, with examples of pre- and post-synaptic marker apposition highlighted by arrows. **b-d**, GENtoniK increases density of SYN1, PSD-95, and their apposition expressed as punctate per neurite length (n = 16 wells from n = 2 independent experiments). **eh**, GENtoniK promotes excitability and mature resting properties in day 28 hPSC-cortical neurons. **e,** >90% of treated neurons fired evoked action potentials in contrast to <40% of DMSO controls. Traces show representative responses for each group. **f-h**, Quantification of electrophysiology parameter AP frequency (**f**), AP threshold (**g**), and resting membrane potential (**h**) (n = 11 neurons per group from 4-6 dishes and 3 independent experiments). **i-m,** RNA-seq and CUT&RUN (3 biological replicates) reveal that GENtoniK induces shift from immature to mature transcriptional programs. **i,** Gene ontology analysis showing enrichment for mature neuron function in genes upregulated by the cocktail; and enrichment for immature function and transcriptional regulation in genes downregulated by the cocktail or occupied by DOT1L-target H3K79 2-methylation. **j,** In the BrainSpan Atlas of the Developing Human Brain (https://www.brainspan.org), genes downregulated by GENtoniK display higher average expression during early development and decrease over time (left), genes upregulated by GENtoniK display an average expression that increases from early development to gestation and after birth (right). Top panels show smoothed means curves with confidence intervals, bottom panels show heatmaps of normalized expression **k-m,** CUT&RUN peak profiles of LSD1 and DOT1L targets H3K4 and H3K79 2-methylation in immature, untreated d7 hPSC-cortical neurons across the whole genome (**k**) and in genes downregulated (**l**) or upregulated (**m**) by GENtoniK in RNA-seq. **b-d** and **f-h**, Two-tailed Welch’s *t*-test; asterisks indicate statistical significance. Mean values are represented by a black line (**b-d**) or a bar graph (**f-h**). Error bars represent S.E.M. Scale bars are 50 μm.

Intrinsic electrophysiological features, such as passive membrane properties and the ability to fire action potentials (APs) are also important indicators of functional neuronal maturation^34^. To assess the effect of the drug cocktail on membrane properties and excitability, we performed whole-cell patch-clamp recordings in cortical neurons at day 28 from plating. Similar to the IEG studies, treatment was withdrawn 7 days before recordings to ensure that differences were maturation-mediated and not a direct effect of the ion channel activators NMDA and Bay K 8644. Over 90% of GENtoniK-treated neurons displayed evoked APs compared to less than 40% of control neurons (Fig. 3e). Among AP-firing neurons, those treated with GENtoniK displayed higher firing frequencies (Fig. 3f) and lower AP thresholds (Fig. 3g). Despite resting membrane potential values being significantly more mature in treated neurons (Fig. 3h), their range was still distant from the physiological range of −60 to −70mV reported for the cortex *in vivo*^35^. These results indicate that GENtoniK significantly promotes synaptic connectivity and excitability, but additional, likely extrinsic factors may be required to achieve more mature resting membrane properties.

### GENtoniK induces immature to mature shift in transcription

We next conducted RNA sequencing to assess global changes in gene expression induced by the smallmolecule treatment. In accordance with a dual effect of the cocktail on chromatin state and calcium influx, we treated hPSC-cortical neurons with either the two epigenetic factors, the two calcium channel agonists, or the complete GENtoniK cocktail (Supp. Fig. 5a). Genes differentially expressed in GENtoniK were similarly regulated by the epigenetic drugs alone but to a lesser magnitude, which is consistent with the hypothesis that calcium influx potentiates transcriptional changes facilitated by chromatin remodeling (Supp. Fig. 5b-d). Although both calcium-channel agonists were identified as maturation enhancers in our protein-based screen, their combined effect on gene expression was modest 7 days after treatment withdrawal (Supp. Fig. 5b).

Gene ontology analyses of transcripts downregulated by GENtoniK revealed enrichment in immature, early post-mitotic neuron functions, including migration and axon guidance, as well as transcriptional regulation (Fig. 3i, Supp. Fig. 5e). Upregulated genes were enriched in mature neuron functionality, including chemical synaptic transmission and transmembrane ion transport (Fig. 3i, Supp. Fig. 5f). While previous studies indicate a switch from glycolytic to oxidative metabolism in maturing neurons^36^, we observed enrichment in both glycolysis and oxidative phosphorylation, as well as fatty acid metabolism in treated cells (Supp. Fig. 6). To match the transcriptional data with chronological changes of gene expression *in vivo*, we plotted differentially expressed genes against the BrainSpan Atlas of the Developing Human Brain dataset^37^. Genes that are downregulated by GENtoniK were more highly expressed in the early embryo and decreased towards birth (Fig 3j, left panel). In contrast, genes upregulated by the treatment generally showed an increase in expression through gestation (Fig 3j, right panel).

We next performed CUT&RUN chromatin profiling on histone marks downstream of the epigenetic factors targeted by the cocktail (Fig. 3k). Although LSD1 can switch its substrate to H3K9 in the mature neuron-specific variant, we focused on its canonical target H3K4 reasoning that maturation-enhancing inhibition likely targets the immature form. In untreated, day 7 cortical neurons, both H3K4 and H3K79 2-methylation were more highly enriched at GENtoniK-downregulated versus GENtoniK-upregulated genes (Fig. 3l, m). H3K4me2 was widespread in the genome, with highest enrichment in the promoter region and near the transcription start site (Supp. Fig. 7a). In contrast, H3K79me2 was enriched at a much smaller subset of genes, where it extended into the transcribed region (Supp. Fig. 7b). Interestingly, genes within H3K79 peaks showed near-identical ontology enrichment to those downregulated by GENtoniK by RNA-seq, being overrepresented in neuron migration, chromatin modifying, and RNA processing gene categories (Fig. 3i, Supp. Fig. 7c-e). Chromatin regulating genes within H3K79me2 peaks include GENtoniK target LSD1 (Supp. Fig. 7d), while mRNA processing genes with H3K79me2 peaks, such as *NOVA2* and *CELF1* (Supp. Fig. 7e), have been shown to participate in cortical neuron development^38,39^. These results indicate that H3K79 methylation may play a role in maintaining immature gene expression programs, and that loss of this mark might facilitate neuronal maturation in GENtoniK-treated cells.

### GENtoniK enhances maturation across neuronal culture systems

We next tested the efficacy of GENtoniK across hPSC-derived neuronal systems. Because our screen relied on the female hESC line H9 (WA09), we first replicated the results in male cortical neurons and derived from induced pluripotent stem cell (iPSCs) lines, confirming GENtoniK’s effect on maturation across different hPSC lines (hESC versus hiPSC) and across both sexes (Supp. Fig. 8).

Alternative maturation strategies are routinely employed in neuronal cultures, including the addition of trophic factors such as brain-derived neurotrophic factor (BDNF) and the use of culture media with more physiological levels of glucose and ion concentrations (BrainPhys)^40^. We conducted time course experiments to assess efficacy and compatibility of GENtoniK with existing maturation approaches. GENtoniK in standard Neurobasal medium (without neurotrophic factors) induced neuronal maturation parameters more robustly than the combination of both BrainPhys and BDNF, while treatment with GENtoniK in combination with BrainPhys and neurotrophic factors showed an additional, albeit modest increase in maturation (Supp. Fig. 9).

Self-organizing 3D culture systems such as brain organoids have become a widely used model system to study human brain development and disease^41^. However, similar to 2D culture systems, 3D organoids are subject to slow maturation rates^42^. We observed that forebrain organoids treated with GENtoniK from day 15-50 of derivation, displayed an increased density of SYN1 puncta (Fig 4a, b), and increased number of cells with nuclear expression of EGR1 and FOS (Fig. 4c, d, Supp. Fig. 10) at day 60. For these studies, organoids were not subjected to KCl stimulation before IEG immunostaining, thus indicating higher levels of spontaneous activity following GENtoniK treatment. GENtoniK-treated organoids also displayed lower expression of immature neuron marker DCX (Supp. Fig 10).

**Fig. 4.**
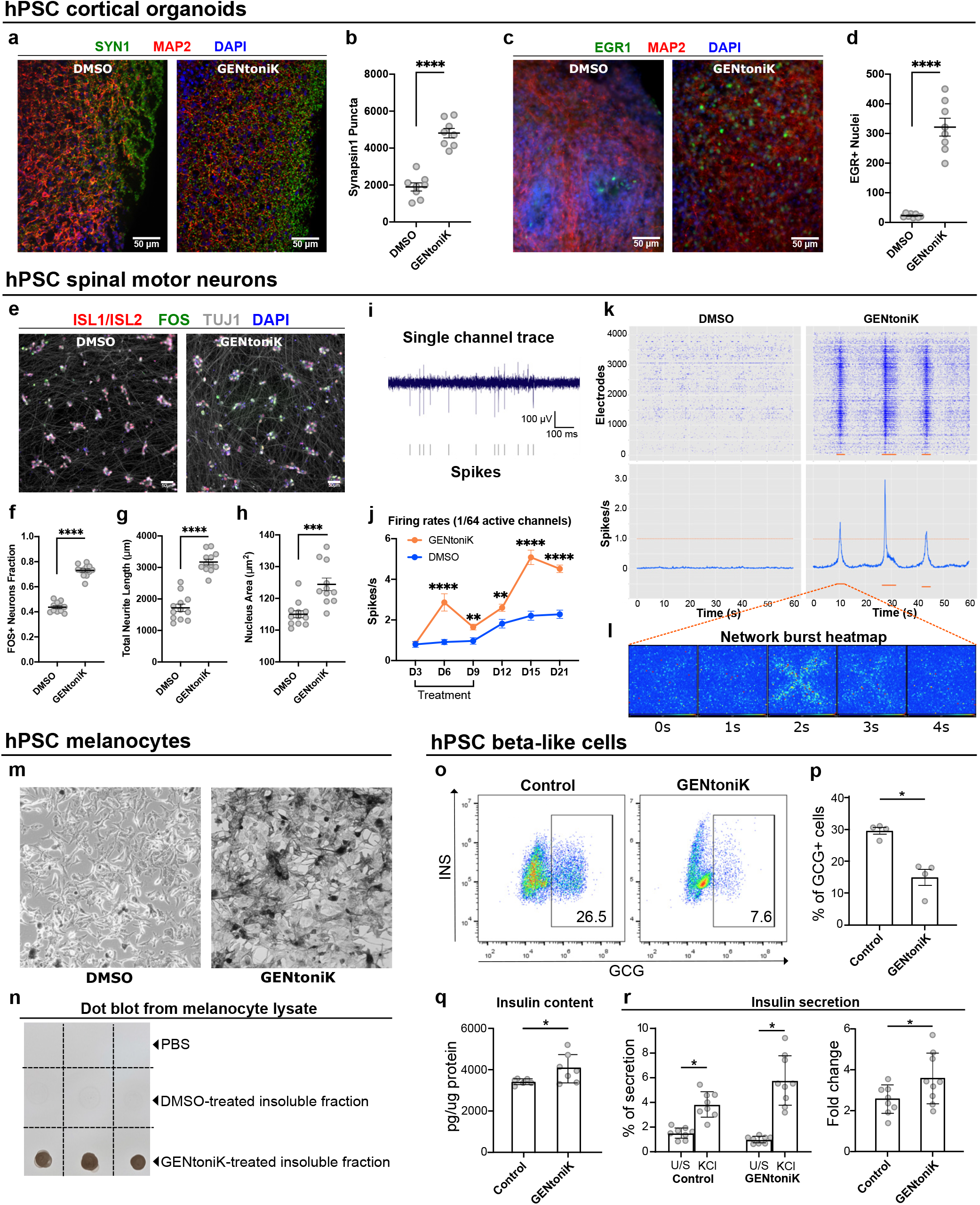
Validation of maturation strategy across neuronal and non-neuronal hPSC-derived cells. **a-d,** GENtoniK treatment induces synaptogenesis and spontaneous activity in cortical organoids. **a,** Representative images of immunofluorescent staining for SYN1 and MAP2 in day 60 organoids. **b**, Quantification of total SYN1 puncta per field (n = 8 cryosections randomly sampled from n = 20 organoids). **c,** Representative images of immunofluorescence staining for EGR1 and MAP2 in unstimulated day 60 organoids. **d,** Quantification of EGR1+ cells per field (n = 8 cryosections randomly sampled from n = 20 organoids). **e-h,** GENtoniK promotes maturation of hPSC-derived spinal motor neurons. **e,** Representative high-content maturation assay images of ISL1/2+ spinal motor neurons (day 40 of hPSC differentiation). **f-h**, Quantification showing GENtoniK-improved KCl-induction of FOS+ cells (**f**), total neurite length (**g**), and nucleus area (**h**) in SMNs (n = 12 microplate wells from 2 independent differentiations). **i-l**, GENtoniK treatment increases firing rates and induces spontaneous bursting activity on SMNs plated on high-density multielectrode arrays. **i**, Sample single channel trace of GENtoniK-treated SMNs illustrating spike detection. **j**, Time-course analysis of average firing rates in SMNs plated on HD-MEAs, calculated from 60 s of activity in the 1/64 most active electrodes (n = 128 electrodes from 2 MEA probes). **k**, Representative 60-second spike rastergrams (top) and average firing rates (bottom) of SMNs plated on a HD-MEAs. Only GENtoniK-treated SMNs displayed spontaneous bursting events (orange bars). **l**, Whole array heatmap of a 4-second bursting event. **m-n,** GENtoniK treatment induces early pigmentation in hPSC-melanocytes. **m**, Brightfield images of melanocytes (day 33 of hPSC differentiation) that received GENtoniK or DMSO from day 11. **n**, Dot blot analysis of PBS or cell extract of melanocytes treated with GENtoniK or DMSO (n = 3 biological replicates). **o-r**, GENtoniK promotes maturation of hESC-derived beta-like cells. Representative flow cytometry analysis (**o**) and quantification (**p**) of the percentage of GCG+ cells in INS-GFP+ cells after 7 days treatment with GENtoniK or control followed by 2 days treatment-free (n = 4 biological replicates). **q**, Total insulin content of INS-GFP+ cells after 7 days treatment with GENtoniK or control followed by 2 days treatment-free (n = 6-7 biological replicates). **r**, Static KCl-stimulated human insulin secretion and fold change in beta-like cells after 7 days treatment with GENtoniK or control followed by 2 days treatment-free. The assay was performed in the presence of 2 mM D-glucose (n = 8-9 biological replicates). **b**, **d, f-h, j** and **p-r**, Two-tailed Welch’s *t*-test; asterisks indicate statistical significance. Mean values are represented by a black line (**b**, **d**, **f-h**) or a bar graph (**p-r**). Error represent S.E.M. Scale bars are 50 μm.

We next addressed whether the treatment can drive the maturation of hPSC-derived neurons outside the cortex or forebrain. ISL1+ spinal motor neurons (SMNs) treated with GENtoniK displayed a highly significant increase across all the maturity parameters tested (Fig. 4e-h). We observed that SMNs exhibit high levels of spontaneous activity when cultured on high-density multielectrode arrays (Fig. 4i). In a timecourse experiment, average firing rates were increased modestly in the presence of the drug cocktail (possibly via direct ion channel activation effect). In contrast, a more pronounced effect was observed starting 6 days after treatment withdrawal indicating that the treatment triggered a long-lasting maturation effect (Fig 4j). Intriguingly, only SMNs pretreated with GENtoniK exhibited highly synchronous bursts of activity in the 0.8-0.6 Hz range (Fig. 4k, l), reminiscent of spontaneous network activity episodes observed in the embryonic spinal cord^43^.

### GENtoniK enhances cell function in non-neuronal lineages

Slow maturation rates of human PSC-derived cells are a common problem across lineages beyond neurons. To assess the potential of GENtoniK in other cell types, we next looked at neural crest-derived melanocytes which produce the pigment melanin in a maturation-dependent manner. The production and secretion of melanin from melanocytes is responsible for human skin and hair color, and hPSCs-melanocytes have been used to model various pigmentation disorders^44^. Using our established differentiation protocol^45^, treatment of hPSC-derived melanocytes with GENtoniK, starting at day 11, induced a dramatic increase in pigmentation at day 33 of differentiation, compared to untreated melanocytes (Fig. 4m, n).

Finally, we tested GENtoniK on a cell type derived from a different germ layer, hPSC-derived insulin-secreting pancreatic beta cells. These cells arise from definitive endoderm^46^ and are of great interest in the development of cell-based treatments for type I diabetes^47^. Although many protocols have been reported, one major limitation is the generation of a subset of glucagon(GCG)+insulin(INS)+ polyhormonal cells^48^. Flow cytometry analysis revealed that GENtoniK treatment decreased the number of GCG+ cells among INS+ cells (Fig. 4o, p). Importantly, beta-like cells that received GENtoniK treatment from days 20 to 27 of differentiation displayed evidence of improved functional maturation including increased total insulin content, fraction of insulin granules, and KCl-induced insulin secretion at day 29 (Fig. 4q-r; Supp. Fig. 11). These results suggest that GENtoniK can trigger some aspects of cell function and maturation even in non-neural lineages.

## Discussion

We present a combined chemical strategy aimed at promoting the maturation of human stem cell-derived neurons, which we obtained by combining hits from a high-content small molecule screen. Applying a multiparameter readout enabled us to identify compounds that effectively drive neuronal maturation rather than simply promoting individual features such as neurite outgrowth^49,50^. PCA of the screen results yielded three phenotypic clusters of compounds that either promoted or inhibited neuronal maturation and compounds that promoted the growth of non-neural contaminants. The enrichment of mTOR and PI3K regulators among maturation inhibitors concurs with recent findings proposing mTOR activation as driver of interneuron maturation^51^. An unexpected finding was the identification of TGF-β and ROCK-inhibitors as compounds promoting a “flat cell” non-neuronal fate, which is a known contaminant of neural differentiations and thought to represent a neural crest^52^ or fibroblast-derived^53^ mesenchymal cell lineage. Both TGF-β and ROCK-inhibitors are commonly used across many neural differentiation protocols, but our results indicate that they may promote undesired cell types if used at later differentiation stages.

A central finding of our study was the presence of an epigenetic program in immature neurons that prevents rapid maturation of human neurons. We hypothesize that GENtoniK acts in a two-pronged manner. The epigenetic probes GSK2879552 and EPZ-5676 induce a shift in chromatin accessibility from an immature (migration, axon guidance) to a mature transcriptional program (synaptic transmission, ion channel subunits). We further speculate that those changes in chromatin state facilitate NMDA and Bay K 8644-mediated activation of calcium-dependent transcription^28^ as an additional driver of maturation.

We identified several inhibitors of LSD1 in our primary screen. LSD1 has been reported to regulate differentiation and maturation in olfactory^25^ and cortical neurons^54,55^, specifically as a member of the CoREST repressor complex. In addition to its roles in development, LSD1 participates in a myriad of functions in a highly context- and complex-specific manner^56^, highlighting the importance of limiting the time of treatment to avoid off-target effects. Alternatively, functional specificity could be mediated by targeting individual complexes. Although a CoREST-specific probe has been developed^57^, in our hands it was highly toxic to neuronal cultures preventing an assessment of any direct effects on maturation (data not shown). DOT1L has been shown to modulate cell-cycle exit during neuronal differentiation^58^, but its role in regulating post-mitotic maturation has not been studied. Our chromatin profiling data in immature neurons indicate that DOT1L substrate H3K79me2 could be involved in controlling the accessibility of other transcriptional regulators including LSD1, making it an intriguing candidate as a potential master regulator of gene expression during development. In agreement with this observation, H3K79me2 levels have been shown to increase globally alongside chromatin condensation during neuronal differentiation^59^, suggesting it might participate in establishing an “epigenetic barrier” during the transition from pluripotent cells to neural progenitors and immature neurons; a barrier then retained in human neurons for protracted periods during neuronal maturation. The apparent absence of a demethylase for this mark make it a plausible timekeeper, as its valence appears to be determined by the rate of nucleosome turnover^60^.

We demonstrate that the same chemical strategy promotes aspects of functional maturation in non-neuronal cells, but more in-depth studies will be required to define the optimal formulation to drive maturation across other cell types. For example, while NMDA receptors and voltage-gated calcium channels have demonstrated functions in melanocytes and pancreatic beta cells^61–64^, their activation might be dispensable to drive maturation in other cells, where alternative factors such as hormones might be required instead. Similarly, there may be alternative epigenetic regulators that contribute to maturation rates across distinct cell types and organ systems to assure appropriate tissue and species-specific timing of maturation. Recent studies have shown that differences in the rate of biochemical reactions including protein synthesis and degradation correlate with species-specific differences in somite and spinal cord development^65,66^. However, further studies are needed to demonstrate a causal relationship and to elucidate whether those mechanisms also apply to later developmental stages such as neuronal maturation. GENtoniK provides a simple, alternative, and likely complementary strategy to accelerate the timing of maturation in neuronal and some non-neural cell types. Furthermore, the use of GENtoniK may facilitate the application of human PSC technology in capturing more mature, adult-like states in modeling human development and disease.

## Methods

### Cell Culture

<u>Human pluripotent stem cells (hPSCs)</u>, both embryonic and induced, were maintained in Essential 8 medium (Thermo) on Vitronectin-coated plates as previously described^67^. Cells were passaged twice per week and collected for differentiations within passages 30 to 50. Mycoplasma testing was conducted every 2 months.

<u>hPSC-derived excitatory cortical neurons</u> were generated using a protocol based on the previously described dual-SMAD inhibition paradigm^68^. Briefly, hESC were dissociated into single cells with Accutase and seeded at 250,000/cm^2^ onto Matrigel-coated plates in Essential 8 medium with 10 μM Y-27632. During days 1 to 10 of the protocol, medium consisted of Essential 6 (Thermo) with 10 μM SB431542 (Tocris) and 100 nM LDN193189 (Stemgent). Wnt inhibitor XAV-939 at 2 μM was included from day 1 to 3 to improve anterior patterning^69^. On days 11-20, medium consisted of N2-supplemented DMEM/F12 (Thermo). Cells received daily medium exchanges throughout the differentiation. On day 20 cells were dissociated in Accutase for 30 m and cryopreserved in STEM-CELLBANKER solution (Amsbio) at 10 million cells/ vial. Neurons were thawed as needed for experiments and plated on poly-L-ornithine and laminin-coated plates (PLO/Lam), in low-glucose (5 mM) Neurobasal-A medium supplemented with 2% B27 and 1% GlutaMAX (Thermo). Neurons received medium exchanges twice per week. During the first 7 days after plating, medium was supplemented with notch-inhibitor DAPT at 10 μM to force lingering progenitors out of the cell cycle^70^.

<u>Primary embryonic rat cortical neurons</u> (Thermo) were thawed following vendor instructions and maintained in the same manner as hPSC-cortical neurons.

<u>Spinal motor neurons</u> derivation was adapted from a previously described protocol^71^ to feeder-free monolayer culture. In brief, Accutase-dissociated hESCs were seeded at 600,000/cm^2^ onto Geltrex-coated plates and underwent dual-SMAD inhibition in the presence of CHIR99021 and Smoothened agonist. On day 11, spinal progenitors were collected and plated on poly-d-lysine, laminin, and fibronectin-coated (PDL/Lam/FN) plates and maintained in N2/B27 medium containing Smoothened agonist, retinoic acid, BDNF, GDNF, CTNF, and DAPT. On day 24, SMNs were re-plated on PDL/Lam/FN and maintained in Neurobasal medium supplemented with 2% B-27, ascorbic acid, retinoic acid, BDNF, GDNF, and CTNF. Treatment with GENtoniK or DMSO was initiated the day after re-plating.

<u>Dorsal forebrain organoid</u> generation was adapted from a previously reported protocol^72^. Briefly, 10,000 EDTA-dissociated hPSCs were plated per well of a 96-well V-bottom low-attachment plate (S-bio). Cells were allowed to self-aggregate in hPSC growth medium overnight. From days 1 to 8, medium was changed every two days with Essential 6 supplemented with 10 μM SB431542, 100nM LDN193189, and 2 μM XAV-939. On day 8, media was switched to organoid growth medium consisting of a 50:50 mixture of Neurobasal and DMEM/F12 with 1% NeuroBrew 21 (Miltenyi), 0.5% N2, 1% GlutaMAX, 0.5% MEM non-essential amino acids solution, 0.1% 2-mercaptoethanol, and 1μM recombinant human insulin (Sigma). Organoids were collected from the wells on day 14 and transferred to 10cm dishes at roughly 20 organoids per dish. Dishes were placed on an orbital shaker set to gentle motion to prevent organoid fusion.

<u>Melanocyte</u> differentiation was executed as previously reported^73^. In brief, the day before differentiation, hPSCs with were plated on Matrigel at 200,000 cells per cm^2^ in E8 medium with 10μM Y-27632. From days 0 to 11 of the protocol, cells received daily exchanges of Essential 6 containing: 1ng/ml BMP4, 10μM SB431542 and 600nM CHIR99021 (days 0-2); 10μM SB431542 and 1.5μM CHIR99021 (days 2-4); 1.5μM CHIR99021 (days 4-6); and 1.5μM CHIR99021, 5ng/ml BMP4 and 100nM EDN3 (days 6-11). On day 11, melanoblasts were sorted using a BD-FACS Aria6 cell sorter at the Flow Cytometry Core Facility of MSKCC. Cells were dissociated into single cells with Accutase for 20 minutes and then stained with an APC-conjugated antibody against cKIT (Invitrogen). Cells positive for APC (cKIT) were sorted and 4, 6-diamidino-2-phenylindole (DAPI) was used to exclude dead cells. Upon FACS sorting, cKIT+ melanoblasts were plated onto dried PO/Lam/FN dishes. Cells were fed with melanocyte medium every 2 to 3 days and passaged using Accutase at a ratio of 1:4 once a week. Melanocyte media consisted of Neurobasal supplemented with: 50ng/ml SCF, 500 μM cAMP, 10ng/ml FGF2, 3 μM CHIR99021, 25ng/ml BMP4, 100nM EDN3, 1mM L-glutamine, 0.1 mM MEM NEAA, 2% B27 and + 2% N2.

<u>Pancreatic beta cell</u> differentiation was performed using *INS^GFP/W^* MEL-1 cells. Cells were cultured on Matrigel-coated 6-well plates in StemFlex medium (Thermo Fisher) and maintained at 37°C with 5% CO2. MEL-1 cells were differentiated using a previously reported strategy^74^. Briefly, on day 0, cells were exposed to basal medium RPMI 1640 (Corning) supplemented with 1× GlutaMAX (Thermo Fisher), 50 μg/mL Normocin, 100 ng/mL Activin A (R&D systems), and 3 μM of CHIR99021 (Cayman Chemical) for 24 hours. The medium was changed on day 2 to basal RPMI 1640 medium supplemented with 1× GlutaMAX, 50 μg/mL Normocin, 0.2% FBS (Corning), 100 ng/mL Activin A for 2 days. On day 4, the resulting definitive endoderm cells were cultured in MCDB131 medium supplemented with 1.5 g/L sodium bicarbonate, 1 ×glutamax, 10 mM glucose, 2% BSA, 50 ng/ml FGF7, 0.25 mM ascorbic acid for 2 days. On day 6, the cells were differentiated in MCDB131 medium supplemented with 2.5 g/L sodium bicarbonate, 1× GlutaMAX, 10 mM glucose, 2% BSA, 0.25 mM ascorbic acid, 2 μM retinoic acid, 0.25 μM SANT1, 50 ng/ml FGF7, 200 nM TPB, 200 nM LDN193189 and 0.5× ITS-X supplement for 2 days to pancreatic progenitor stage 1 cells. On day 8, the cells were induced to differentiate to pancreatic progenitor stage 2 cells in MCDB131 medium supplemented with 2.5 g/L sodium bicarbonate, 1×glutamax, 10 mM glucose, 2% BSA, 0.25 mM ascorbic acid, 0.2 μM retinoic acid, 0.25 μM SANT1, 2 ng/ml FGF7, 100 nM TPB, 400 nM LDN193189 and 0.5× ITS-X supplement for 3 days. On day 11, the cells were induced to differentiate to insulin expressing cells in MCDB131 medium supplemented with 1.5 g/L sodium bicarbonate, 1×glutamax, 20mM glucose, 2% BSA, 0.1 μM retinoic acid, 0.25 μM SANT1, 200 nM LDN193189, 1 μM T3, 10 μM ALKi5, 10 μM zinc sulfate, 10 μg/mL heparin and 0.5×ITS-X for 3 days. On day 14, the cells for static or dynamic KCl stimulated insulin secretion (KSIS) analysis were scraped off from plates and relocated onto 24mm insert and 3.0 μm polycarbonate membrane, 6-well tissue culture trans-well plate into hemispherical colonies and the cells for insulin content analysis and flow cytometry analysis were kept on original plates. All the cells then were further maturated in MCDB131 medium supplemented with 1.5 g/L sodium bicarbonate, 1×glutamax, 20 mM glucose, 2% BSA, 100 nM LDN193189, 1 μM T3, 10 μM zinc sulfate, 10 μg/mL heparin, 100 nM GS in XX and 0.5× ITS-X for 7 days. Then cells were further matured in MCDB131 medium supplemented with 1.5 g/L sodium bicarbonate, 1×glutamax, 20 mM glucose, 2% BSA, 1 μM T3, 10 μM zinc sulfate, 10 μg/mL heparin, 1 mM acetylcysteine, 10 μM Trolox, 2 μM R428 and 0.5× ITS-X with GENtoniK or control treatment for 7 days.

### Small molecule treatment

A bioactive compound library containing 2688 compounds was used for screening at a concentration of 5 μM (Selleck Bioactive Library, Selleck Chemicals). 192 DMSO wells contained within the library were used as negative controls. For confirmation of primary hits, compounds were extracted from the library plates with an Agilent Bravo liquid handling platform and re-subjected to the high-content assay in triplicates at 5 μM. 22 confirmed compounds were purchased from Selleck Chemicals, reconstituted in a suitable solvent and applied for dose-response validation in a concentration log scale (30nM, 100nM, 300nM, 1000nM, 3000nM, 10,000 nM). GENtoniK cocktail was defined as a mixture of 4 small molecules: GSK2879552, EPZ-5676, Bay K 8644, and NMDA, applied at a working concentration of 1 μM each. Stocks of individual GENtoniK ingredients were reconstituted in DMSO to 10mM (GSK2879552, EPZ-5676, Bay K 8644), or in water to 50 mM (NMDA) and stored at −20 C until the day of experiments. Unless stated otherwise, controls received a corresponding volume of DMSO (3:10,000).

### Immunostaining

#### Monolayer cultures

Cells were fixed in 4% paraformaldehyde in PBS for 30 m, permeabilized for 5 m in PBS with 0.1% Triton X-100 and blocked for 30 m in PBS with 5 % normal goat serum (NGS). Incubation with primary antibodies was performed overnight at 4 C at the specified dilution in PBS with 2% NGS. Following 3 washes with PBS, cells were incubated with fluorescently conjugated secondary antibodies (2 μg/ml) for 30 m at room temperature. Nuclear staining with DAPI at 1 μg/ml was simultaneous to secondary antibody incubation. For high-content experiments, all steps were assisted by automated liquid handling at the MSKCC Gene Editing and Screening Core Facility. A list of antibodies used in this study is presented in Supplementary Table 2.

#### Forebrain organoids

Organoids were collected in 1.5 ml centrifuge tubes, washed in PBS, and fixed with 4% paraformaldehyde solution in PBS overnight at 4 C. Fixed organoids were rinsed in PBS and equilibrated in a solution of 30% weight/volume sucrose in PBS for 24 h or until sunk to the bottom of the tube. Organoids were embedded in OCT compound (Fisher) on cryomolds, frozen and sectioned to a thickness of 30 μm in a cryostat. Sections were collected in 1 ml centrifuge tubes (1 per antibody), washed in TBS with 0.3% Triton-X and blocked in the same solution with 10% NGS. Primary antibody incubation was done overnight in TBS with 0.5% Tween-20, and followed by washes, and secondary antibody incubation for 2 h at RT in the same buffer. Sections were mounted on slides with ProLong medium (Fisher) and imaged on a Zeiss microscope equipped with a 20x high numerical aperture objective and an Apotome optical sectioning system (Zeiss). For quantification of SYN1 puncta, images were batch-analyzed using the Synapse Counter ImageJ plugin^75^.

### High-content imaging

#### High-content maturity assay

Cortical neurons were seeded PLO/Lam-coated 384-well plates at a density of 5000/well and maintained as described. For bioactive compound screening, compounds were added 7 days after plating to a final concentration of 5 μM in replicate plates. Following 7 days of treatment, cells were rinsed twice and maintained in plain medium for an additional 7 days. Before fixation, one replicate plate was stimulated with 50 mM KCl for 2 hours. Immunostaining for FOS, EGR1, and MAP2 and counterstaining with DAPI was performed as described above. Images (4 fields/well at 20x magnification) were captured through an INCell Analyzer 6000 HCA system (GE Healthcare).

#### Image analysis and quantification of screen results

Phenotypic analysis of screen images was conducted using the Columbus software (Perkin Elmer). Extracted parameters included total number of nuclei, nuclear area, nuclear roundness index (DAPI); total neurite length per nucleus (MAP2); and fraction of FOS-positive, EGR1-positive and double-IEG positive nuclei (FOS/EGR1). For IEG quantification, ratios of positive nuclei were calculated by applying a threshold of fluorescence intensity within DAPI-positive nuclei. IEG nuclei ratios in unstimulated plates were then subtracted from KCl-stimulated plates to isolate the KCl depolarization-mediated response. Morphological variables (nuclear and neurite) were averaged between unstimulated and KCl plates. Sequential b-score and z-score normalization and principal component analysis were performed in the KNIME analytics platform^76^ with the High Content Screening Tools extension.

#### Synaptic marker analysis

hPSC-cortical neurons were thawed and plated on PLO/Lam 96-well plates. Drug treatment was initiated after 7 days and maintained for 21 day. Cells were fixed after an additional 7 days in plain medium. Immunostaining for Synapsin 1, PSD95, and MAP2 was conducted as described above. 10 images per well were captured using the confocal modality of the IN Cell 6000 HCA system. A mask was applied to the area surrounding MAP2-positive processes, and SYN1 and PSD95 puncta were quantified within the defined region. For quantification of pre- and post-synaptic marker apposition, a mask was applied to an area containing and immediately surrounding SYN1 puncta, and PSD95 puncta localized within this region were counted. Synaptic puncta counts per field were normalized to total neurite length.

### Electrophysiology

#### Whole-cell patch-clamp

hPSC-cortical neurons were plated onto PLO/Lam-coated 35mm dishes at a density of 75k/cm^2^. Treatment with GENtoniK or DMSO began 7 days after plating and maintained for 14 days. Recordings were initiated 7 days after treatment withdrawal, within days 28 to 33 from plating. Whole-cell recordings were performed at 23 – 24° C while the cells were perfused in freshly made ACSF containing (in mM): 125 NaCl, 2.5 KCl, 1.2 NaH_2_PO_4_, 1 MgSO_4_, 2 CaCl_2_, 25 NaHCO_3_ and 10 D-glucose. Solutions were pH-corrected to 7.4 and 300-310 mOsm. Neurons were recorded with pipettes of 3-7 MΩ resistance filled with a solution containing (in mM): 130 potassium-gluconate, 4 KCl, 0.3 EGTA, 10 Na_2_-phosphocreatine, 10 HEPES, 4 Mg_2_-ATP, 0.3 Na_2_-GTP and 13 biocytin, pH adjusted to 7.3 with KOH and osmolarity to 285–290 mOsmol/kg. Recordings were performed on a computer-controlled amplifier (MultiClamp 700B Axon Instruments, Foster City, CA) and acquired with an AxoScope 1550B (Axon Instruments) at a sampling rate of 10 kHz and low-pass filtered at 1 kHz.

#### Multi-electrode array recording

hPSC-derived spinal motor neurons were seeded onto poly-l-lysine-coated complementary metal oxide semiconductor multi-electrode array (CMOS-MEA) probes (3Brain)^77^. A 100-μl droplet of medium containing 200,000 neurons was placed on the recording area. After 1 h incubation, 1.5 ml of medium were added to the probe and replaced every 3 days. Cells received treatment with GENtoniK or DMSO during days 3 to 9 from plating. Recordings were performed every 3 days for 18 days, 24 h after medium changes. 1 minute of spontaneous activity was sampled from 4096 electrodes using the BioCAM system and analyzed using BrainWave 4 software. Spikes were detected using a sliding window algorithm on the raw channel traces applying a threshold for detection of 9 standard deviations. Network bursts were detected by applying a hard threshold of 1 spike/second on the entire 4096-channel array.

### Gene expression and chromatin profiling

#### RNA-seq

RNA was extracted using the Direct-zol RNA miniprep kit (Zymo). Total RNA samples were submitted to GENEWIZ for paired-end sequencing at 30-40 million reads. Analysis was conducted in the Galaxy platform^78^. Transcript quantification was performed directly from adapter-trimmed FASTQ files using the Salmon quasi-mapping tool^79^ referenced to GENCODE Release 36 (GRCh38.p13) transcripts. DESeq2^80^ was used for differential expression analysis from Salmon-generated transcript per million (TPM) values. Differentially expressed genes with a Benjamini-Hochberg adjusted p-value below 0.05 and a baseMean cutoff of 1000 were applied to gene set overrepresentation analysis using the Goseq tool^81^. For gene set enrichment, all genes with a baseMean above 1000 were analyzed using the GSEA software^82^.

#### CUT&RUN

hPSC-derived cortical neurons were collected 7 days after plating for CUT&RUN chromatin profiling using the standard protocol^83^. Antibodies against H3K4me2 (Upstate), H3K79me2 (Active Motif) and mouse IgG (Abcam) were used at 1:100 for 100k cells per antibody. DNA was collected via phenolchloroform extraction and submitted to the MSKCC Integrated Genomics Operation core for paired-end sequencing at 5 million reads. Analysis was performed in the Galaxy platform. Following alignment to ENSEMBL GRCh38 genome build using Bowtie 2^84^, peaks were called using MACS^85^, and visualized with ChIPSeeker^86^ and deepTool2^87^, using mouse IgG as control for normalization.

### Dot blot for melanocyte pigmentation

hESC-melanocytes were dissociated in Accutase, rinsed, and collected in PBS. A pellet containing 1M cells was lysed in 50 μl RIPA buffer with sonication, and centrifuged at 10,000 RCF for 3 m. After discarding the supernatant, the insoluble fraction was resuspended in 80 μl of PBS. 10 μl of this solution was applied to a nitrocellulose membrane, air dried, and imaged with a standard office scanner to assess pigmentation.

### Pancreatic beta cell maturation assays

#### Flow cytometry analysis

hESC-derived cells were dissociated using Accutase, fixed and permeabilized using Fixation/Permeabilization Solution Kit (BD Biosciences) according to the manufacturer’s instructions. Briefly, cells were first fixed with fixation/permeabilization buffer for 30 mins at 4°C in dark and then washed twice with washing buffer with 10 mins incubation each time at room temperature. Then, the fixed cells were incubated with primary antibody overnight at 4°C, washed twice with washing buffer with 10 mins incubation each time at RT. After 30 mins incubation with fluorescence-conjugated secondary antibody at 4°C, cells were washed twice with washing buffer with 10 mins incubation each time at room temperature and re-suspended in PBS buffer for analysis. The following primary antibodies were used: anti-Insulin (1:50, Dako) and anti-Glucagon (1:100, Abcam). Samples were analyzed with an Accuri C6 flow cytometry instrument and the data were processed using FlowJo v10 software.

#### Static and dynamic KSIS

On day 30 cells were starved in 2 mL glucose-free pancreatic beta cells maturation media and followed by 2 mL glucose-free DMEM (with GlutaMAX) for 1 hour and additional 1 hour incubation in KRBH buffer (containing 140 mM NaCl, 3.6 mM KCl, 0.5 mM NaH_2_PO_4_, 0.2 mM MgSO_4_, 1.5mM CaCl_2_, 10 mM Hepes (pH 7.4), 2 mM NaHCO_3_ and 0.1% BSA) in a 5% CO_2_/37°C incubator. To perform static KSIS, cells were exposed sequentially to 100 μL of KRBH with 2 mM glucose, or 2 mM glucose with 30 mM KCl; supernatants were collected after 60 mins and spun down to eliminate the cells and debris. Supernatants were used for ELISA (Insulin Chemiluminescence ELISA Jumbo, Alpco). To measure the total insulin levels in cells in each sample, cells were lysed in RIPA buffer supplemented with 1×protease inhibitor cocktail (ThermoFisher Scientific) with vortexing for 2 mins at RT and flash freeze the samples in liquid nitrogen and thaw to help the lysis and release the cellular insulin. Lysates were spun down, and supernatant was used for ELISA. Insulin secretion from cells in each condition was normalized to KRBH treatment. To perform dynamic KSIS, cells were embedded in chambers with the order of filter paper-biogel P4 beads-cells-biogel P4 beads order sandwich and then the chambers were installed on the biorep perfusion system (Biorep Technology) and first perfused with Krebs buffer containing 2 mM glucose at a flow rate of 100 μL/min and followed by perfusion with 2 mM glucose + 30 mM KCl for 25 mins. Insulin secretion from cells in each fraction in KCl stimulation were normalized to KRBH treatment.

#### Insulin content measurement

D30 hESC-derived beta-like cells were dissociated using Accutase and resuspended in DMEM containing 2% FBS and 1 mM EDTA. 80,000 *INS*-GFP^+^DAPI^-^ cells were FACS sorted by an ARIA2 instrument, washed once with PBS and lysed in 200 μL RIPA buffer supplemented with 1 × protease inhibitor cocktail (ThermoFisher Scientific). The insulin content was measured by ELISA.

#### Immuno-electron microscopy

To analyze granular ultrastructure, control or chemical treated-hPSC-derived beta-like cell clusters were washed with serum-free media and fixed with 2.5% glutaraldehyde, 4% paraformaldehyde, 0.02 % picric acid in 0.1 M buffer. After three buffer washes, the cell clusters were fixed again using 1% OsO_4_^-^1.5%K-ferricyanide at RT for 60 mins followed by three buffer washes. After dehydration steps of 50%, 70%, 85%, 95%, 100%, 100%,100% EtOH, the cell clusters were infiltrated with 100% EtOH mixed 1:1 with acetonitrile, followed by acetonitrile, acetonitrile 1:1 with EMbed 812 epoxy resin, resin and finally, embedded in fresh resin which was polymerized at 50 deg C for 36 hr. Sections were cut at 65 nm and picked up on nickel grids. Sections were washed with saturated Na-periodate, followed by 50 mM glycine, and blocking buffer. Then, the sections were stained with anti-insulin antibody at original dilution followed by 10 nm gold Goat anti-Guinea pig IgG (Aurion, 1:100). Samples were imaged with a JEOL JEM 1400 TEM with an Olympus-SIS 2K x 2K Veleta CCD camera.

### Statistical analysis

Averages are reported as arithmetic means +/- SEM (standard error of the mean) unless otherwise indicated. Statistical significance was marked by asterisk notation as follows: (ns) p > 0.05, (*) p ≤ 0.05, (**) p ≤ 0.01, (***) p ≤ 0.001, (****) p ≤ 0.0001. Biological replicates are defined as independent differentiations of a given hPSC line unless indicated otherwise.

## Data Availability

Data generated during this study are deposited at NCBI GEO under accession numbers GSE172544 (RNA-seq) and GSE172543 (CUT&RUN).

## Acknowledgements

We would like to thank members of the Studer lab for continuous support and insightful discussions. We are grateful to M. Fennell for guidance in the conception and design of the screening assay. We thank J. Muller of the MSK Light Microscopy Instrument Cluster and Y. Lin of the Vierbuchen lab for their help in live-imaging experiments. In addition, we would like to thank A. Maccione and M. Falappa of 3Brain AG for assisting in the interpretation of multielectrode array results. We also thank Dr. Lee Cohen-Gould and Mr. Juan Pablo Jimenez at Weill Cornell Medicine Microscopy and Image Analysis Core Facility for help with immunoelectron microscopy. The work was supported in part by a grant for the Tri-institutional stem cell initative from the Starr Foundation and by R01AG054720 to LS, NYSTEM DOH01-STEM5-2016-00300-C32599GG to L.S and S.C, and the core grant P30CA008748. A.B. was supported by a Swiss National Science Foundation Postdoc.mobility fellowship P400PB_180672.; A.P.M. was supported by fellowship F31AG067709-01; R.M.W. was supported by an F32 Ruth L. Kirschstein Postdoctoral fellowship (MH116590); JL is supported by the Rohr Family Research Scholar Award and an Irma T. Hirschl and Monique Weill-Caulier Awar; G.C was supported by an EMBO long-term postdoctoral fellowship and a NYSTEM postdoctoral fellowship.

## Author contributions

E.H.: Conception, study design, data analysis and interpretation, development and execution of high-content assays, hPSC maintenance and differentiation, bioinformatics, multielectrode array recording, writing of manuscript. Y.Z.: design, optimization, and execution of high-content assays, data interpretation. Z.Z.: beta cell differentiation, insulin content and secretion assays, flow cytometry, data analysis, writing of manuscript H.M.: single-cell electrophysiology and data analysis. E.L.C.: development and execution of spinal motor neuron derivation protocol. A.B.: optimization and execution of melanocyte differentiation protocol. A.P.M.: cortical organoid generation and maintenance. R.M.W.: development and supervision of organoid generation. C.L & J.L.: design and supervision of single cell electrophysiology. R.G.: design and supervision of high-content assays. S.C.: data analysis and interpretation, writing of manuscript, G.C.: conception, development of cortical differentiation protocol, L.S.: conception, study design, data analysis and interpretation, writing of manuscript.

## Legends to Supplementary Figures

**Supplementary Figure 1.**
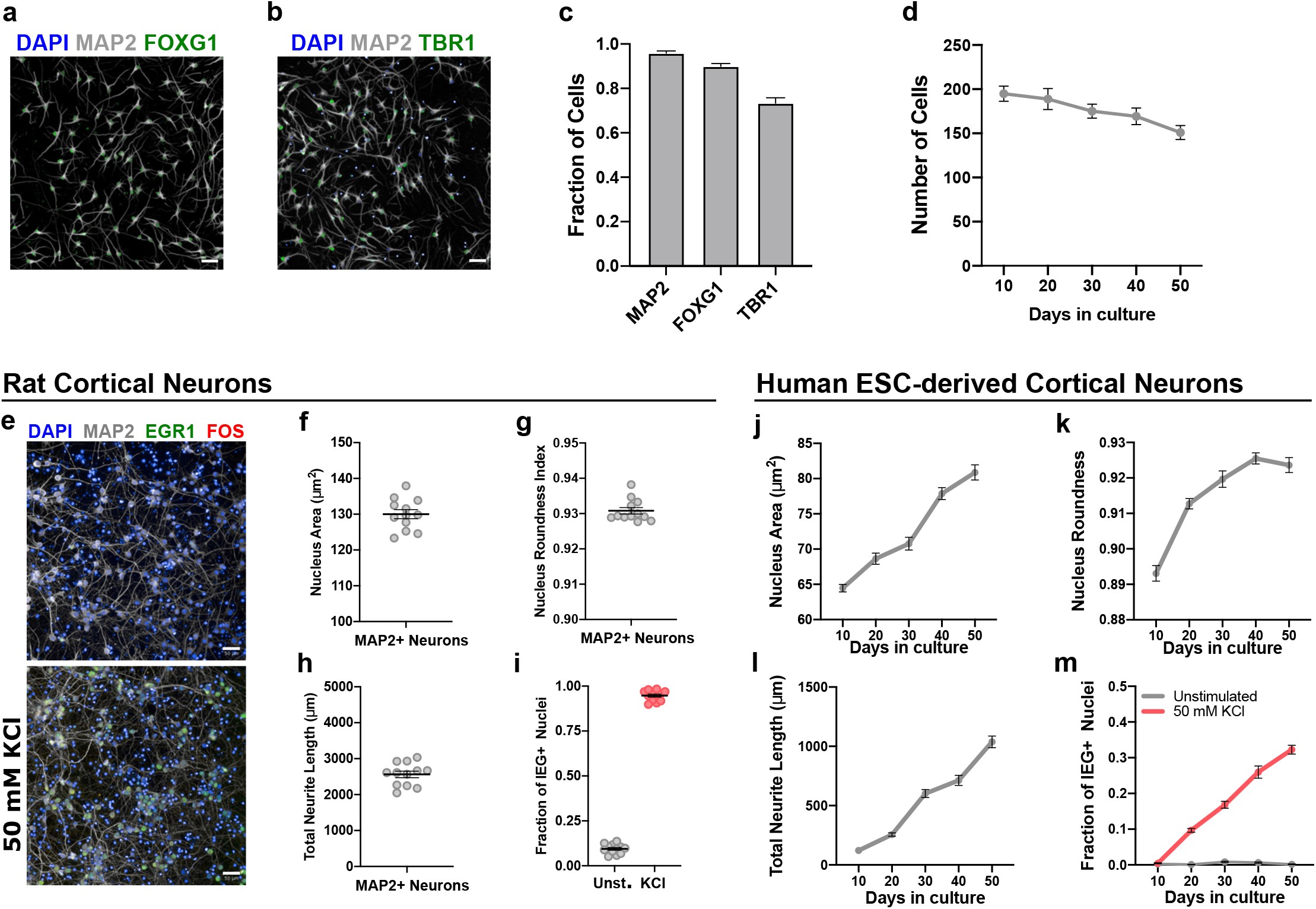
Design and optimization of high-content maturation assay. **a-c**, immunofluorescent staining of day 10 hPSC-cortical neurons for pan-neuronal marker MAP2 (**a,b**), forebrain marker FOXG1 (**a**), and deep-layer cortex marker TBR1 (**b**). **c**, Quantification of immunofluorescent staining (n = 12 microplate wells). **d**, Time-course quantification of cell number in postmitotic hPSC-cortical neurons (DAPI+ cells per field, n = 24 microplate wells). **e**, Immunofluorescent staining of primary embryonic rat cortex neurons (E18) using high-content markers. **f-i**, Quantification of maturation parameters primary rat neurons demonstrate mature values for nucleus size (**f**), nucleus roundness (**g**), neurite length (**h**), and KCl-induced IEG expression (**i**) (n = 12 microplate wells). **j-m**, Time course quantification of maturation parameters in hPSC-derived cortical neurons showing time-dependent increases in nucleus size (**j**), nucleus roundness (**k**), neurite length (**l**), and KCl-induced IEG expression (**m**) (n = 24 microplate wells). Mean values are represented by a black line. Error bars represent S.E.M. Scale bars are 50 μm.

**Supplementary Figure 2.**
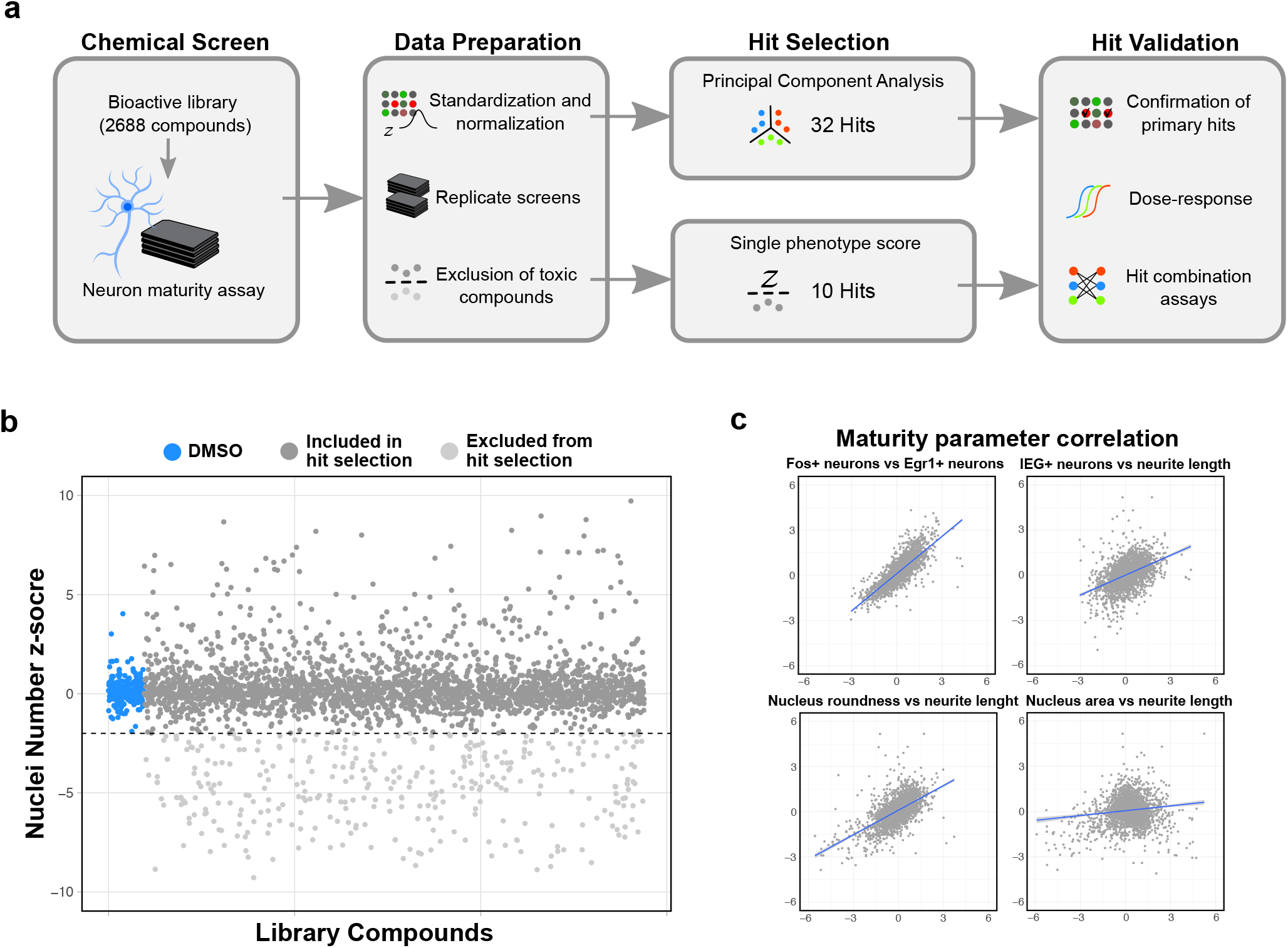
High-content screen data preparation and analysis. **a**, Pipeline of analysis of high-content screen using a 2688-compound bioactive library. Normalization scores (z-scores) of 2 independent screens were averaged and used for selection of hits via PCA or single-parameter scores. **b**, Exclusion of toxic compounds with a mean z-score of total cell number below −2. Note that increases in total cell number were only observed for compounds inducing non-neural cells (Fig. 1e). **c**, Correlation of mean maturation z-scores from 2 screen runs among non-toxic compounds.

**Supplementary Figure 3.**
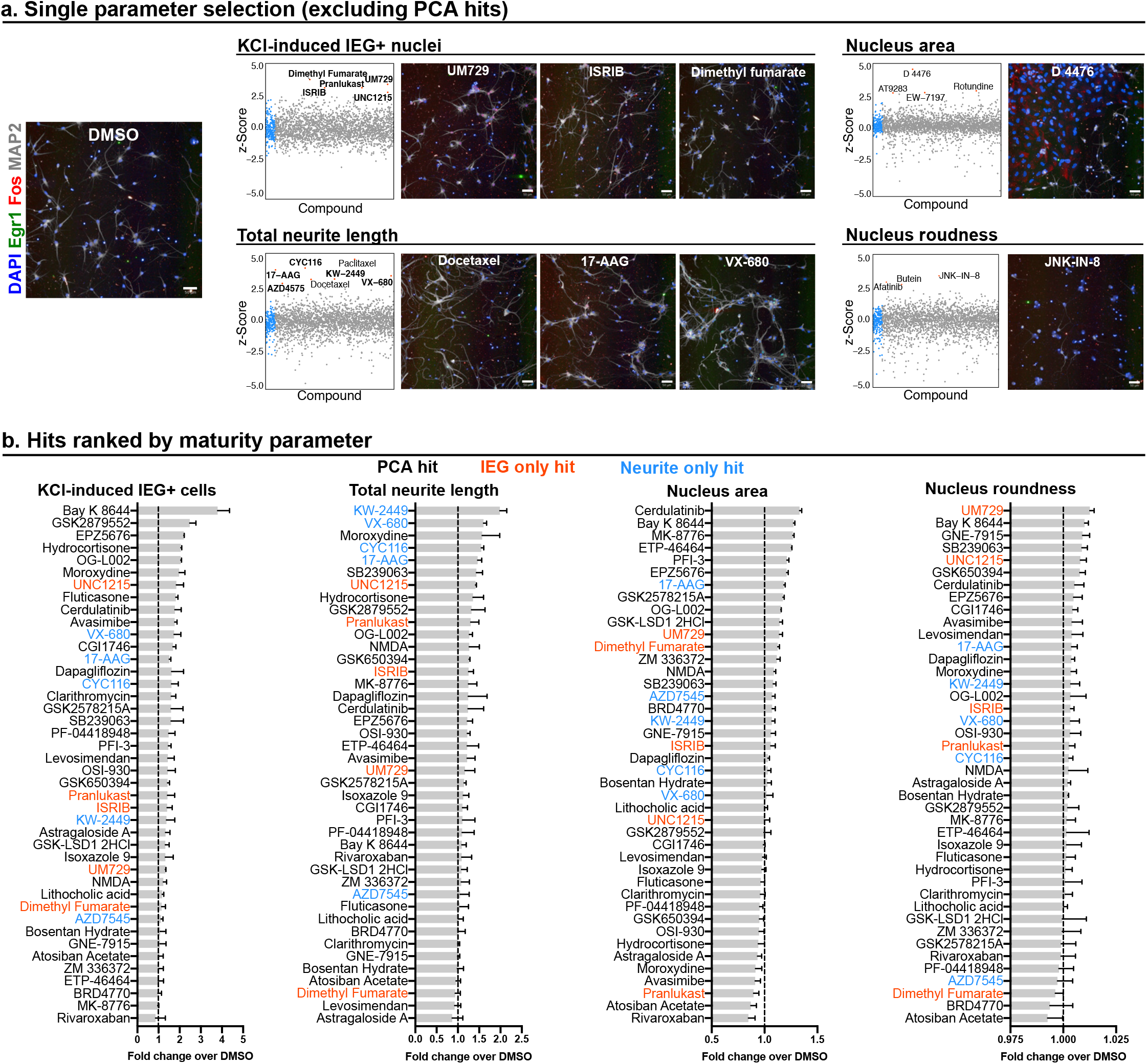
Single parameter hit selection. **a**, Left, representative high-content screen image of a DMSO control well. Right, library compounds (excluding the PCA hits already selected) plotted against individual maturation parameter. Selected compounds are highlighted in **bold**, non-highlighted compounds were not included due to phenotype and/or known molecular target unrelated to neuronal maturation. Screen images are representative of high-scoring compounds for each parameter. **b**, ranking of 42 primary hits (PCA and single parameter) in individual maturation parameters (n = 3 microplate wells). Mean values are represented by bar graph. Error bars represent S.E.M. Scale bars are 50 μm.

**Supplementary Figure 4.**
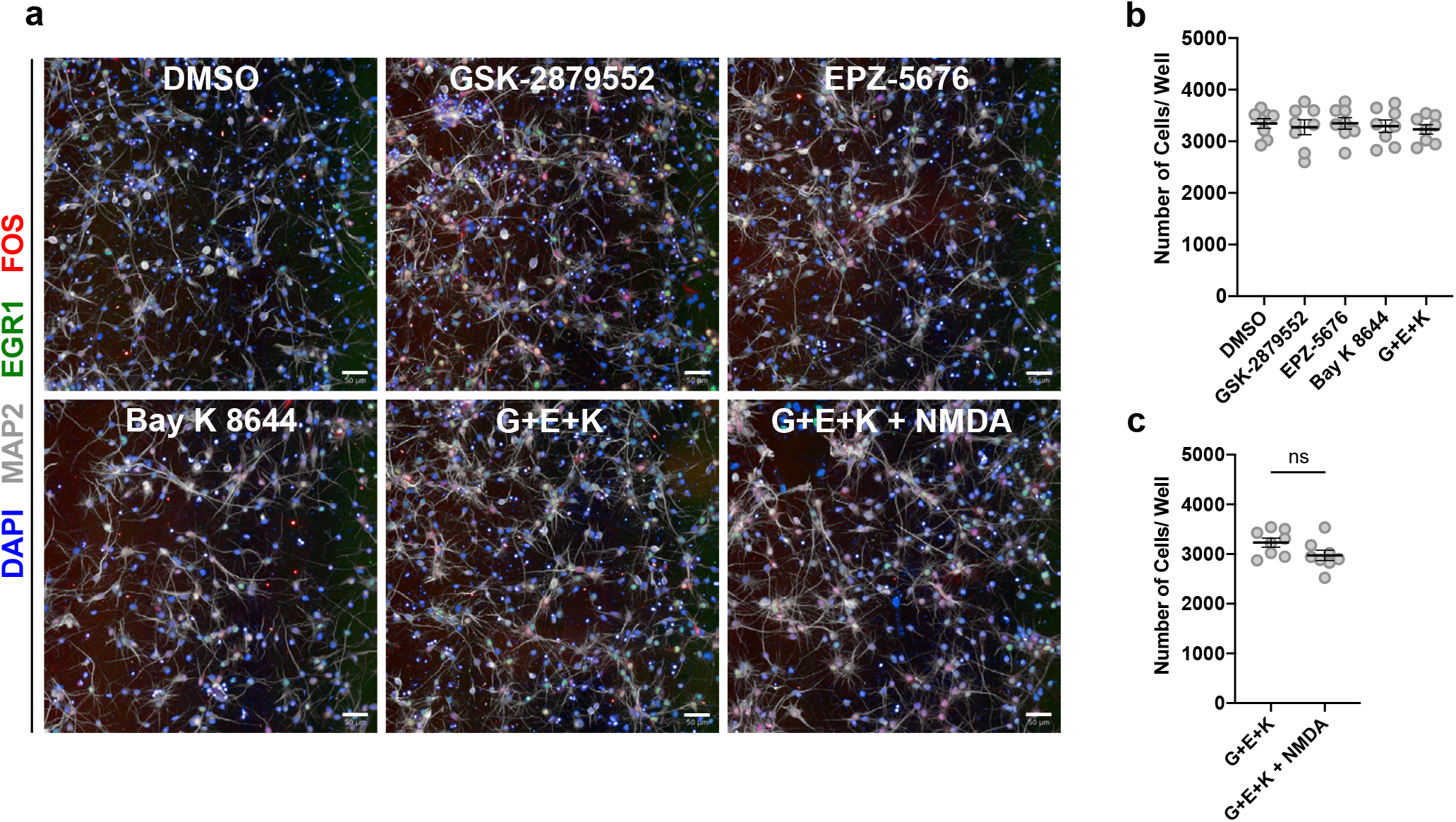
Maturation-promoting small molecules do not significantly affect neuron survival. **a**, Representative staining images from hit combination experiments (**Fig. 2c-e)**, showing day 21 neurons that received the specified treatment from days 7-14. **b**, Quantification of number of cells per well in neurons treated with screen hits GSK-2879552, EPZ-5676, Bay K 8644, and a combination of the 3 (G+E+K) **c**, Quantification of number of cells per well in neurons treated with 3-hit drug combination (G+E+K) and the same with the addition of NMDA. n = 8 microplate wells from 2 independent experiments. Error bars represent S.E.M. Scale bars are 50 μm.

**Supplementary Figure 5.**
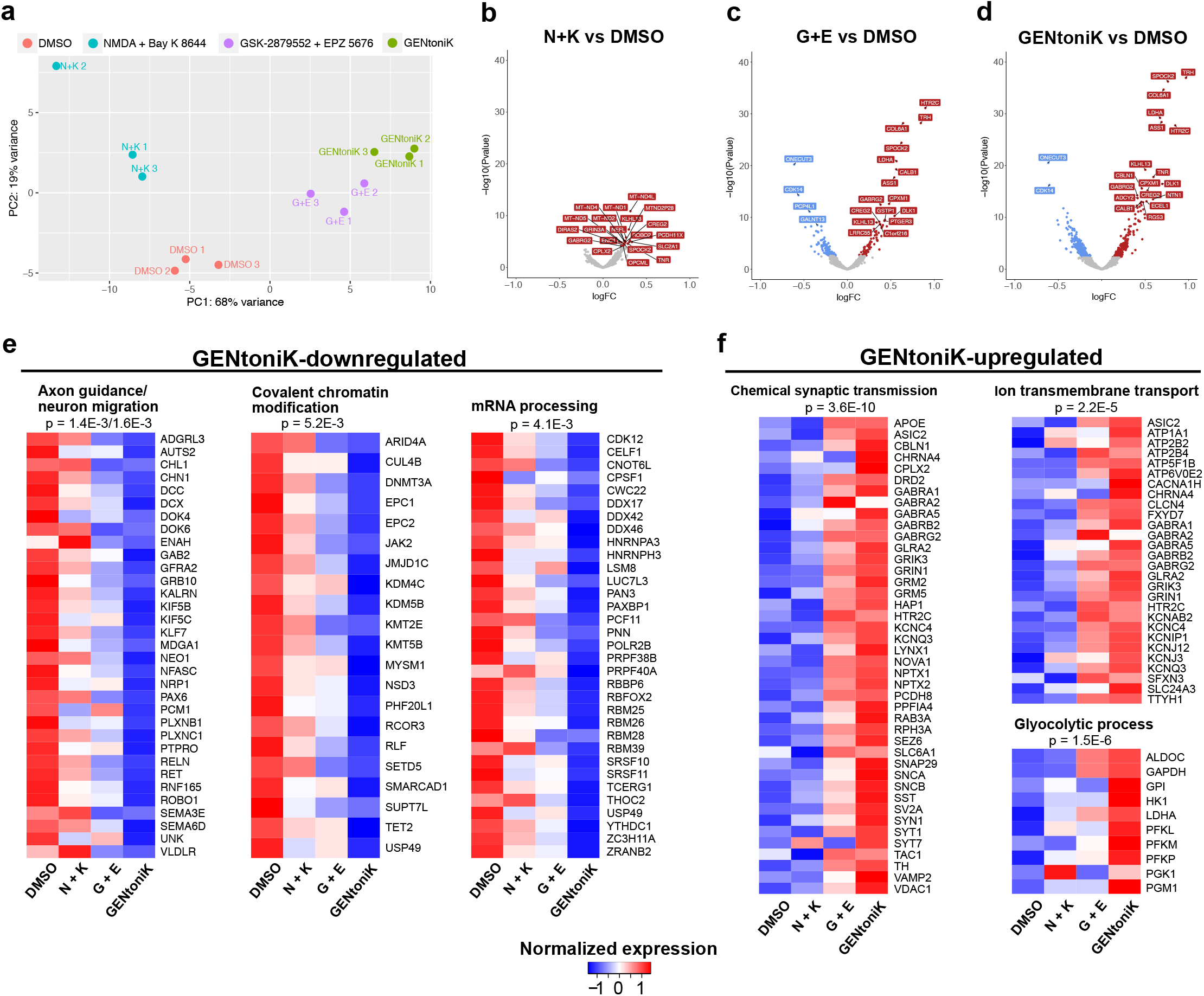
RNA-seq results of day 21 neurons treated with maturation promoting small molecules from d7-14. **a**, Principal component analysis of RNA-seq results from neurons treated with DMSO, two epigenetic drugs (G+E), two calcium influx driving compounds (N+K), or complete GENtoniK. **b-d**, Volcano plots of RNA-seq differential expression analysis vs DMSO of calcium influx agonist NMDA and Bay K 8644 (**b**), epigenetic drugs GSK-2879552 and EPZ-5676 (**c**), or complete GENtoniK (**d**). **e,f,** Heatmaps of genes within overrepresented biological process ontology categories among GENtoniK-downregulated (**e**), and upregulated (**f**) genes. RNA-seq results from 3 biological replicates. Heatmaps show expression normalized by row, calculated from mean TPM values. Displayed p-values are for enrichment of stated gene ontology categories among differentially expressed transcripts.

**Supplementary Figure 6.**
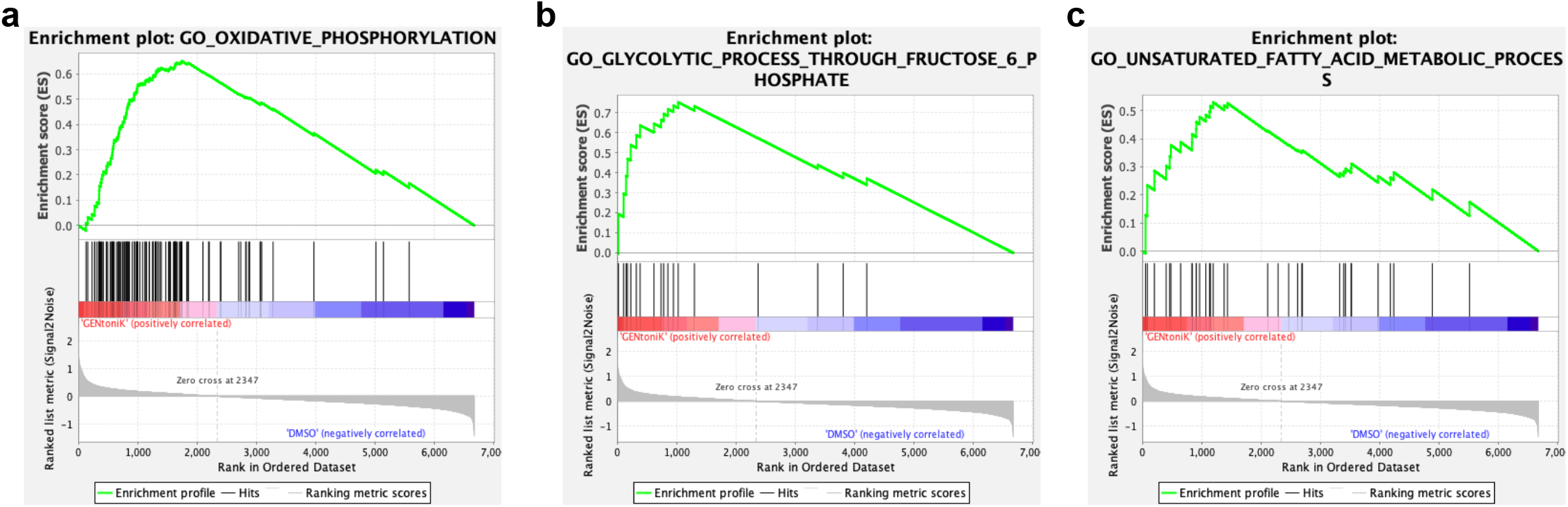
GENtoniK induces transcriptional activation of diverse metabolic pathways in cortical neurons. **a-c,** Gene set enrichment analysis (GSEA) of RNA-seq results showing enrichment for oxidative phosphorylation (**a**), canonical glycolysis (**b**), and fatty acid metabolism (**c**) gene ontology categories enriched in GENtoniK-treated neurons. N = 3 biological replicates.

**Supplementary Figure 7.**
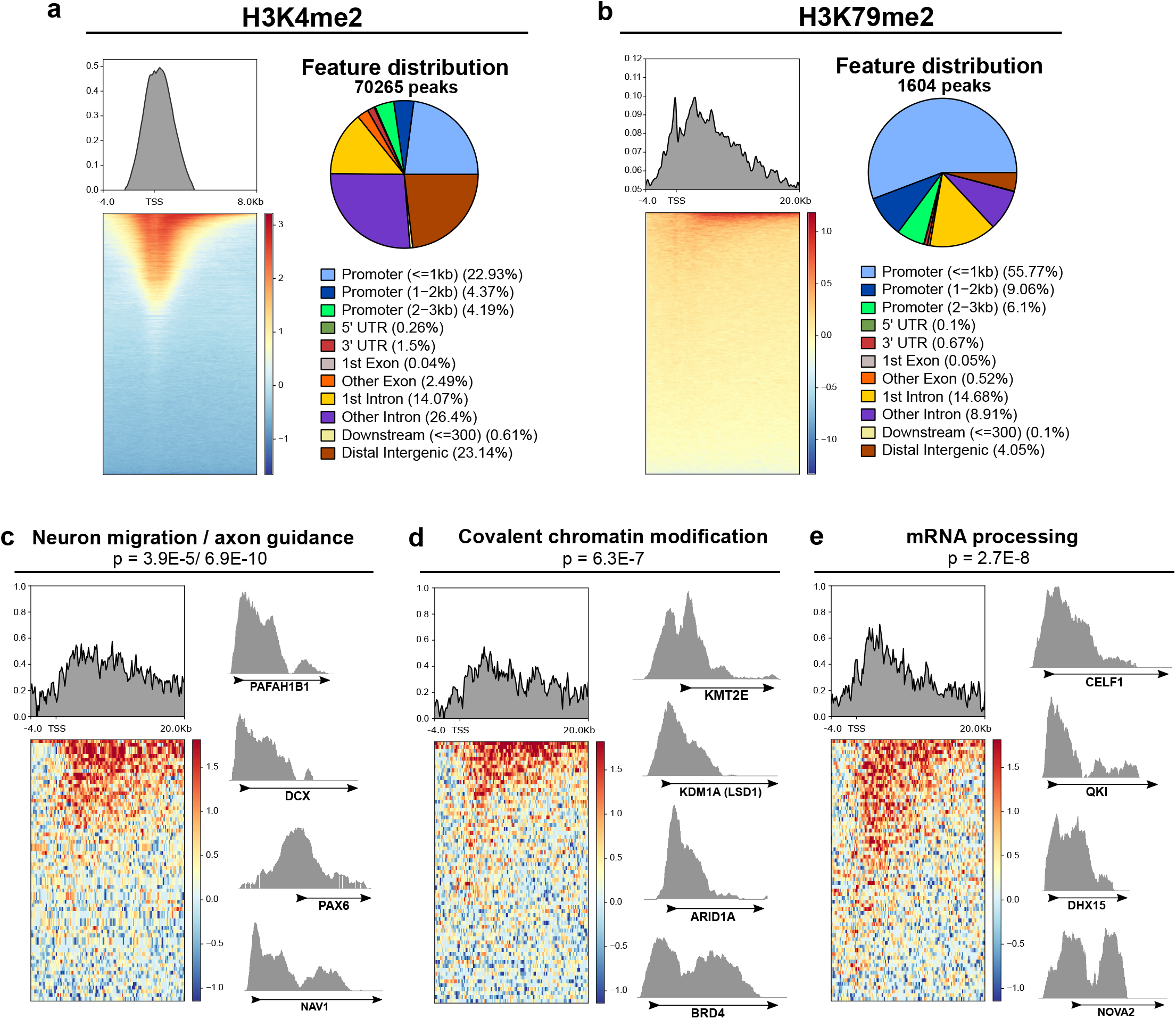
CUT&RUN analysis of LSD1 and DOT1L-targeted histone marks in untreated day 10 immature neurons. **a,** Left, normalized genome enrichment profile of H3K4me2 over IGG control along 12Kb region surrounding the transcription start site (TSS). Right, genome-wide distribution of gene features among H3K4me2 peaks. **b**, Left, normalized genome enrichment profile of H3K79me2 over IGG control along 24Kb region surrounding the transcription start site (TSS). Right, genome-wide distribution of gene features among H3K79me2 peaks. **c-e**, Enrichment of H3K79me2 vs IGG control in gene ontology categories significantly overrepresented among H3K79me2 peaks with representative tracks for genes within each category: GO:0001764-neuron migration and GO:0007411-axon guidance (**c**), GO:0016569-covalent chromatin modification (**d**), and GO:0006397-mRNA processing (**e**). Displayed p-values are for enrichment of stated ontology categories among genes within H3K79me2 peaks.

**Supplementary Figure 8.**
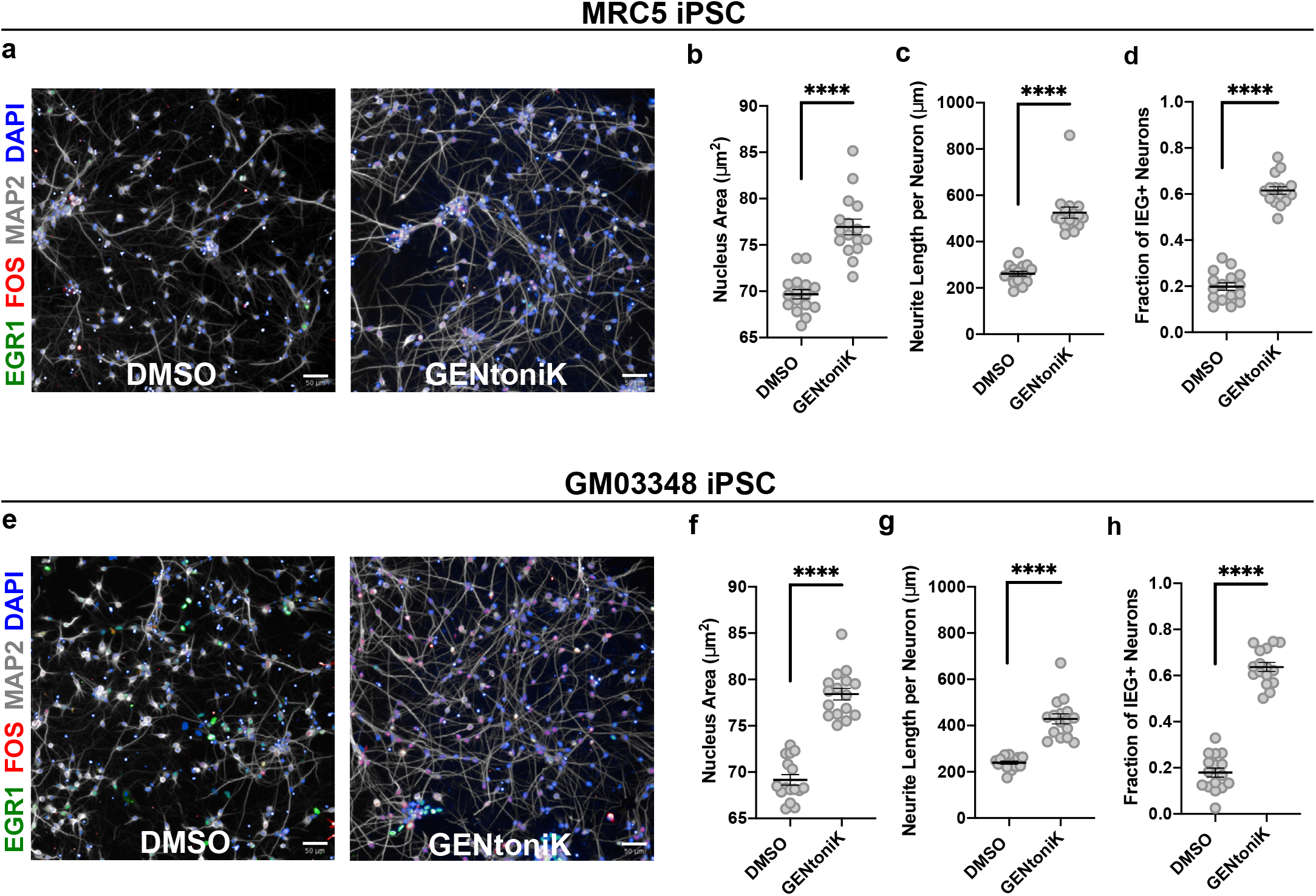
GENtoniK promotes maturation of cortical neurons derived from induced pluripotent stem cells (iPSCs). **a-d,** Neurons derived from reprogrammed normal lung fibroblast line MRC5 (n = 16 microplate wells): representative high-content maturation assay images (**a**), and quantification of maturation parameters nucleus size (**b**), neurite length (**c**) and IEG induction by KCl (**d**). **e-h**, Neurons derived from reprogrammed skin fibroblasts of 10-year-old male (n = 16 microplate wells): representative high-content maturation assay images (**e**), and quantification of maturation parameters nucleus size (**f**), neurite length (**g**) and IEG induction by KCl (**h**). Two-tailed Welch’s *t*-test; asterisks indicate statistical significance. Mean values are represented by a black line. Error bars represent S.E.M. Scale bars are 50 μm.

**Supplementary Figure 9.**
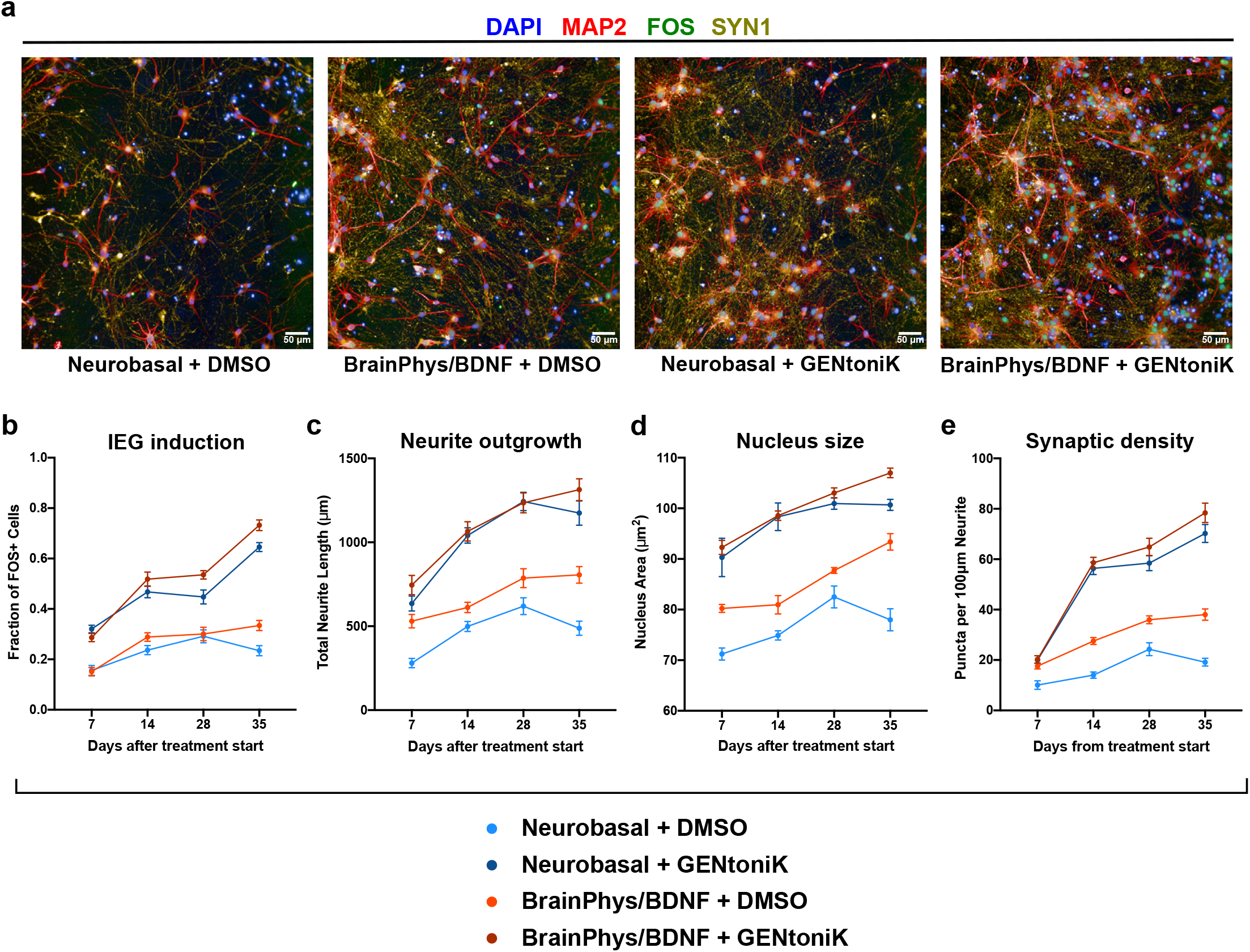
GENtoniK improves upon and complements alternative neuron maturation strategies. **a**, Immunofluorescent stain for MAP2, FOS, and SYN1 of day 35 hPSC-derived cortical neurons in plain Neurobasal medium, BrainPhys medium+BDNF, Neurobasal with GENtoniK, and BrainPhys+BDNF with GENtoniK. **b-e,** Time-course quantification of the maturity parameters: FOS induction by KCl (**b**), neurite length (**c**), nucleus size (**d**), and SYN1 puncta density (**e**) in neurons that received GENtoniK versus DMSO from day 7 from plating. Plates were collected for analysis every 7 days, beginning 7 days after the start of DMSO/GENtoniK treatment. n = 12 microplate wells. Error bars represent S.E.M. Scale bars are 50 μm.

**Supplementary Figure 10.**
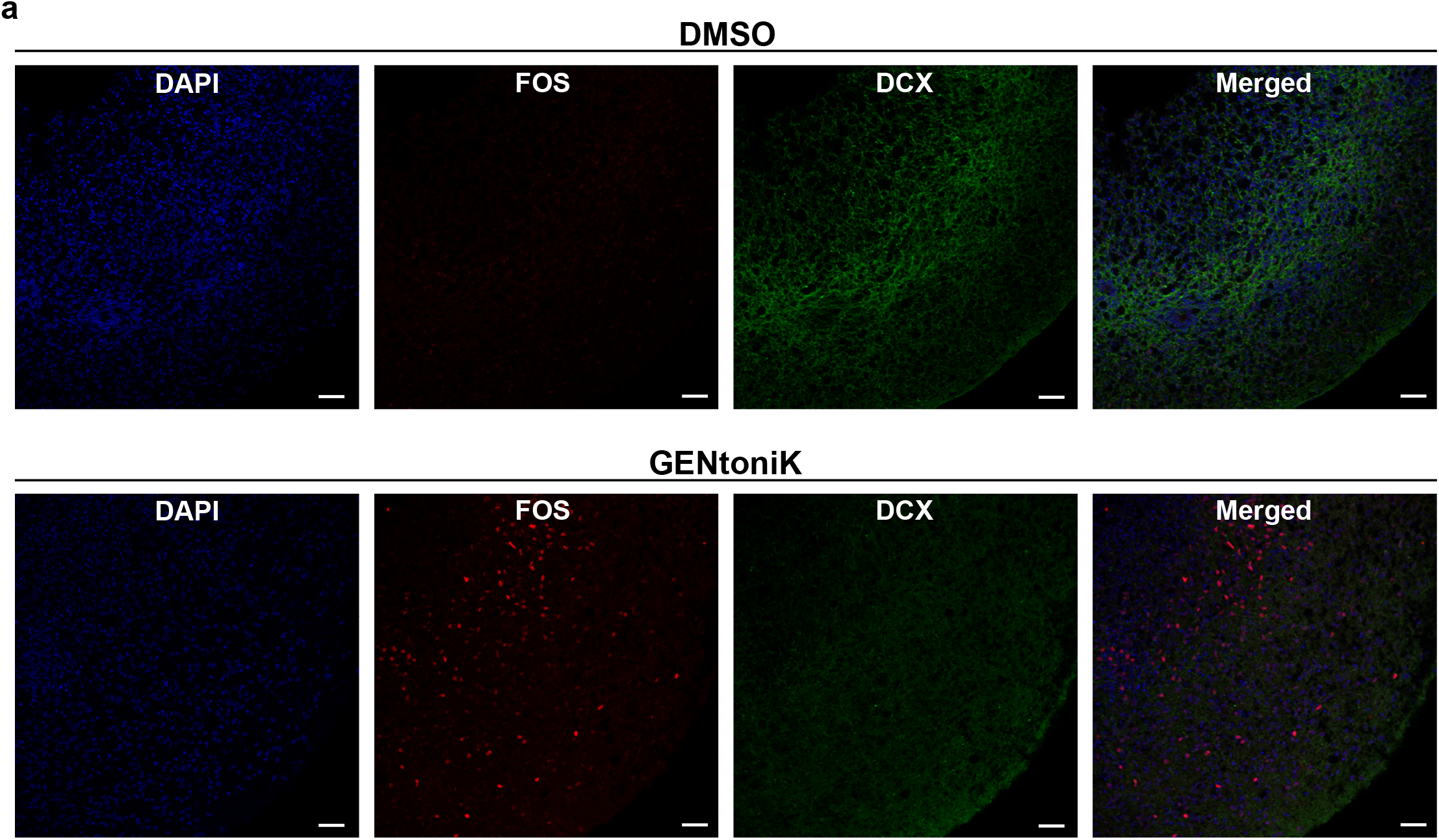
GENtoniK decreases migratory marker expression and increases neuronal activity marker expression in forebrain organoids. **a,** Representative images of immunofluorescent staining for FOS, DCX, and MAP2 in day 60 forebrain organoids that received DMSO (top) or GENtoniK (bottom) from days 15 to 50. Scale bars are 50 μm.

**Supplementary Figure 11.**
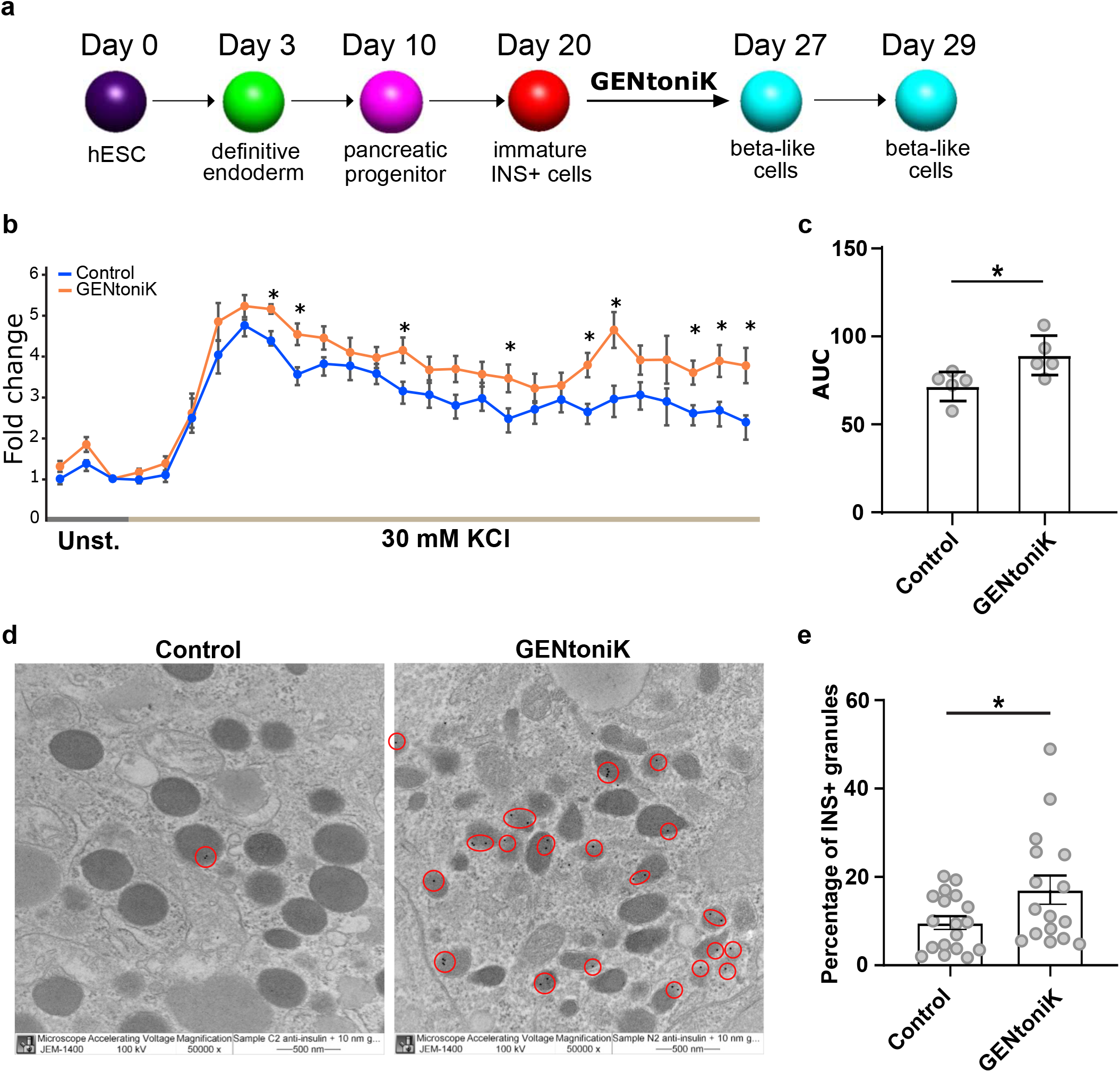
GENtoniK increases dynamic insulin secretion and insulin+ granules in hPSC-derived beta-like cells. **a,** Schematic representation of the stepwise differentiation protocol. hESC-derived immature beta-like cells were treated with GENtoniK or DMSO from days 20 to 27. **b,c,** Dynamic KCl stimulated human insulin secretion (**b**) and area under curve (AUC, **c**) in hESC derived cells after 7 days treatment with GENtoniK or control followed by 2 days treatment-free culture. The assay was performed in the presence of 2 mM D-glucose. Fold change was calculated by dividing the amount of secreted insulin at each time point by the average amount of secreted insulin at 2 mM D-glucose. N = 5 biological replicates. **d**, Representative electron micrographs showing immunogold labelling of insulin in beta-like cells. Circles indicate insulin^+^ granules (10 nm gold particles). Magnification = 50,000x. **e**, Percentage of insulin^+^ granules in control and GENtoniK-treated beta-like cells (N = 16). **c** and **e**, Twotailed Student’s *t*-test; asterisks indicate statistical significance. Error bars represent S.E.M.

## References

1. Kriks, S. et al. Dopamine neurons derived from human ES cells efficiently engraft in animal models of Parkinson’s disease. Nature 480, 547–551 (2011).

2. Shi, Y., Kirwan, P., Smith, J., Robinson, H. P. C. & Livesey, F. J. Human cerebral cortex development from pluripotent stem cells to functional excitatory synapses. Nat. Neurosci. 15, 477–486 (2012).

3. Maroof, A. M. et al. Directed differentiation and functional maturation of cortical interneurons from human embryonic stem cells. Cell Stem Cell 12, 559–572 (2013).

4. Sacai, H. et al. Autism spectrum disorder-like behavior caused by reduced excitatory synaptic transmission in pyramidal neurons of mouse prefrontal cortex. Nat. Commun. 11, (2020).

5. Falke, E. et al. Subicular dendritic arborization in Alzheimer’s disease correlates with neurofibrillary tangle density. Am. J. Pathol. 163, 1615–1621 (2003).

6. Jirsch, J. D. et al. High-frequency oscillations during human focal seizures. Brain 129, 1593–1608 (2006).

7. Ullian, E. M., Sapperstein, S. K., Christopherson, K. S. & Barres, B. A. Control of synapse number by glia. Science (80-.). 291, 657–661 (2001).

8. Benders, M. J. et al. Early brain activity relates to subsequent brain growth in premature infants. Cereb. Cortex 25, 3014–3024 (2015).

9. McAllister, A. K., Katz, L. C. & Lo, D. C. Neurotrophin regulation of cortical dendritic growth requires activity. Neuron 17, 1057–1064 (1996).

10. Barry, C. et al. Species-specific developmental timing is maintained by pluripotent stem cells ex utero. Dev. Biol. 423, 101–110 (2017).

11. Marchetto, M. C. et al. Species-specific maturation profiles of human, chimpanzee and bonobo neural cells. Elife 8, (2019).

12. Linaro, D. et al. Xenotransplanted Human Cortical Neurons Reveal Species-Specific Development and Functional Integration into Mouse Visual Circuits. Neuron 104, 972–986.e6 (2019).

13. Isacson, O. & Deacon, T. Neural transplantation studies reveal the brain’s capacity for continuous reconstruction. Trends Neurosci. 20, 477–482 (1997).

14. Boutros, M., Heigwer, F. & Laufer, C. Microscopy-Based High-Content Screening. Cell vol. 163 1314–1325 (2015).

15. Wu, G. Y., Zou, D. J., Rajan, I. & Cline, H. Dendritic dynamics in vivo change during neuronal maturation. J. Neurosci. 19, 4472–4483 (1999).

16. Ito, K. & Takizawa, T. Nuclear architecture in the nervous system: Development, function, and neurodevelopmental diseases. Frontiers in Genetics vol. 9 (2018).

17. Sheng, M. & Greenberg, M. E. The regulation and function of c-fos and other immediate early genes in the nervous system. Neuron vol. 4 477–485 (1990).

18. Greenberg, M. E. & Ziff, E. B. Stimulation of 3T3 cells induces transcription of the c-fos protooncogene. Nature 311, 433–438 (1984).

19. Murai, J. et al. Chromatin Remodeling and Immediate Early Gene Activation by SLFN11 in Response to Replication Stress. Cell Rep. 30, 4137–4151.e6 (2020).

20. Opitz, T., De Lima, A. D. & Voigt, T. Spontaneous development of synchronous oscillatory activity during maturation of cortical networks in vitro. J. Neurophysiol. 88, 2196–2206 (2002).

21. Singh, S., Carpenter, A. E. & Genovesio, A. Increasing the content of high-content screening: An overview. Journal of Biomolecular Screening vol. 19 640–650 (2014).

22. Shlevkov, E. et al. A High-Content Screen Identifies TPP1 and Aurora B as Regulators of Axonal Mitochondrial Transport. Cell Rep. 28, 3224–3237.e5 (2019).

23. Blazejewski, S., Bennison, S., Liu, X. & Toyo-oka, K. High-Throughput Kinase Inhibitor Screening Reveals Roles for Aurora and Nuak Kinases in Neurite Initiation and Dendritic Branching. bioRxiv 2020.06.25.162271 (2020) doi:10.1101/2020.06.25.162271.

24. Laurent, B. et al. A Specific LSD1/KDM1A Isoform Regulates Neuronal Differentiation through H3K9 Demethylation. Mol. Cell 57, 957–970 (2015).

25. Coleman, J. H., Lin, B. & Schwob, J. E. Dissecting LSD1-Dependent Neuronal Maturation in the Olfactory Epithelium. J. Comp. Neurol. 525, 3391–3413 (2017).

26. Jones, B. et al. The histone H3K79 methyltransferase Dot1L is essential for mammalian development and heterochromatin structure. PLoS Genet. 4, (2008).

27. Kamijo, S. et al. A critical neurodevelopmental role for l-type voltage-gated calcium channels in neurite extension and radial migration. J. Neurosci. 38, 5551–5566 (2018).

28. Bading, H., Ginty, D. D. & Greenberg, M. E. Regulation of gene expression in hippocampal neurons by distinct calcium signaling pathways. Science (80-.). 260, 181–186 (1993).

29. Hou, G. & Zhang, Z. W. NMDA receptors regulate the development of neuronal intrinsic excitability through cell-autonomous mechanisms. Front. Cell. Neurosci. 11, (2017).

30. Rusconi, F. et al. LSD1 modulates stress-evoked transcription of immediate early genes and emotional behavior. Proc. Natl. Acad. Sci. U. S. A. 113, 3651–3656 (2016).

31. Murphy, T. H., Worley, P. F. & Baraban, J. M. L-type voltage-sensitive calcium channels mediate synaptic activation of immediate early genes. Neuron 7, 625–635 (1991).

32. Xia, Z., Dudek, H., Miranti, C. K. & Greenberg, M. E. Calcium influx via the NMDA receptor induces immediate early gene transcription by a MAP kinase/ERK-dependent mechanism. J. Neurosci. 16, 5425–5436 (1996).

33. Liu, X. et al. Extension of cortical synaptic development distinguishes humans from chimpanzees and macaques. Genome Res. 22, 611–622 (2012).

34. Oswald, A. M. M. & Reyes, A. D. Maturation of intrinsic and synaptic properties of layer 2/3 pyramidal neurons in mouse auditory cortex. J. Neurophysiol. 99, 2998–3008 (2008).

35. Dégenètais, E., Thierry, A. M., Glowinski, J. & Gioanni, Y. Electrophysiological properties of pyramidal neurons in the rat prefrontal cortex: An in vivo intracellular recording study. Cereb. Cortex 12, 1–16 (2002).

36. Zheng, X. et al. Metabolic reprogramming during neuronal differentiation from aerobic glycolysis to neuronal oxidative phosphorylation. Elife 5, (2016).

37. Miller, J. A. et al. Transcriptional landscape of the prenatal human brain. Nature 508, 199–206 (2014).

38. Saito, Y. et al. Differential NOVA2-Mediated Splicing in Excitatory and Inhibitory Neurons Regulates Cortical Development and Cerebellar Function. Neuron 101, 707–720.e5 (2019).

39. Popovitchenko, T. et al. Translational derepression of Elavl4 isoforms at their alternative 5’ UTRs determines neuronal development. Nat. Commun. 11, (2020).

40. Bardy, C. et al. Neuronal medium that supports basic synaptic functions and activity of human neurons in vitro. Proc. Natl. Acad. Sci. U. S. A. 112, E2725–E2734 (2015).

41. Chiaradia, I. & Lancaster, M. A. Brain organoids for the study of human neurobiology at the interface of in vitro and in vivo. Nature Neuroscience vol. 23 1496–1508 (2020).

42. Otani, T., Marchetto, M. C., Gage, F. H., Simons, B. D. & Livesey, F. J. 2D and 3D Stem Cell Models of Primate Cortical Development Identify Species-Specific Differences in Progenitor Behavior Contributing to Brain Size. Cell Stem Cell 18, 467–480 (2016).

43. Gonzalez-Islas, C. & Wenner, P. Spontaneous network activity in the embryonic spinal cord regulates AMPAergic and GABAergic synaptic strength. Neuron 49, 563–575 (2006).

44. Mica, Y., Lee, G., Chambers, S. M., Tomishima, M. J. & Studer, L. Modeling Neural Crest Induction, Melanocyte Specification, and Disease-Related Pigmentation Defects in hESCs and Patient-Specific iPSCs. Cell Rep. 3, 1140–1152 (2013).

45. Callahan, S. J., Mica, Y. & Studer, L. Feeder-free derivation of melanocytes from human pluripotent stem cells. J. Vis. Exp. 2016, (2016).

46. Chen, S. et al. A small molecule that directs differentiation of human ESCs into the pancreatic lineage. Nat. Chem. Biol. 5, 258–265 (2009).

47. Mayhew, C. N. & Wells, J. M. Converting human pluripotent stem cells into β-cells: Recent advances and future challenges. Current Opinion in Organ Transplantation vol. 15 54–60 (2010).

48. Teitelman, G., Alpert, S., Polak, J. M., Martinez, A. & Hanahan, D. Precursor cells of mouse endocrine pancreas coexpress insulin, glucagon and the neuronal proteins tyrosine hydroxylase and neuropeptide Y, but not pancreatic polypeptide. Development 118, 1031–1039 (1993).

49. Sherman, S. P. & Bang, A. G. High-throughput screen for compounds that modulate neurite growth of human induced pluripotent stem cell-derived neurons. DMM Dis. Model. Mech. 11, (2018).

50. Sridharan, B. P. et al. A Simple Procedure for Creating Scalable Phenotypic Screening Assays in Human Neurons. Sci. Rep. 9, (2019).

51. Chu, J. et al. Enhanced maturation of human stem cell derived interneurons by mTOR activation. bioRxiv (2019) doi:10.1101/777714.

52. Hu, B. Y. & Zhang, S. C. Directed differentiation of neural-stem cells and subtype-specific neurons from hESCs. Methods Mol. Biol. 636, 123–137 (2010).

53. Tiklová, K. et al. Single cell transcriptomics identifies stem cell-derived graft composition in a model of Parkinson’s disease. Nat. Commun. 11, (2020).

54. Fuentes, P., Cánovas, J., Berndt, F. A., Noctor, S. C. & Kukuljan, M. CoREST/LSD1 control the development of pyramidal cortical neurons. Cereb. Cortex 22, 1431–1441 (2012).

55. Han, X. et al. Destabilizing LSD1 by Jade-2 promotes neurogenesis: An antibraking system in neural development. Mol. Cell 55, 482–494 (2014).

56. Maiques-Diaz, A. & Somervaille, T. C. P. LSD1: Biologic roles and therapeutic targeting. Epigenomics vol. 8 1103–1116 (2016).

57. Kalin, J. H. et al. Targeting the CoREST complex with dual histone deacetylase and demethylase inhibitors. Nat. Commun. 9, (2018).

58. Franz, H. et al. DOT1L promotes progenitor proliferation and primes neuronal layer identity in the developing cerebral cortex. Nucleic Acids Res. 47, 168–183 (2019).

59. Ferrari, F. et al. DOT1L-mediated murine neuronal differentiation associates with H3K79me2 accumulation and preserves SOX2-enhancer accessibility. Nat. Commun. 11, (2020).

60. Chory, E. J. et al. Nucleosome Turnover Regulates Histone Methylation Patterns over the Genome. Mol. Cell 73, 61–72.e3 (2019).

61. Hoogduijn, M. J. et al. Glutamate receptors on human melanocytes regulate the expression of MiTF. Pigment Cell Res. 19, 58–67 (2006).

62. Das, A. et al. Functional expression of voltage-gated calcium channels in human melanoma. Pigment Cell Melanoma Res. 25, 200–212 (2012).

63. Inagaki, N. et al. Expression and role of ionotropic glutamate receptors in pancreatic islet cells. FASEB J. 9, 686–691 (1995).

64. Davalli, A. M. et al. Dihydropyridine-sensitive and -insensitive voltage-operated calcium channels participate in the control of glucose-induced insulin release from human pancreatic β cells. J. Endocrinol. 150, 195–203 (1996).

65. Matsuda, M. et al. Species-specific segmentation clock periods are due to differential biochemical reaction speeds. Science (80-.). 369, (2020).

66. Rayon, T. et al. Species-specific pace of development is associated with differences in protein stability. Science (80-.). 369, (2020).

67. Tchieu, J. et al. A Modular Platform for Differentiation of Human PSCs into All Major Ectodermal Lineages. Cell Stem Cell 21, 399–410.e7 (2017).

68. Chambers, S. M. et al. Highly efficient neural conversion of human ES and iPS cells by dual inhibition of SMAD signaling. Nat. Biotechnol. 27, 275–280 (2009).

69. Tchieu, J. et al. NFIA is a gliogenic switch enabling rapid derivation of functional human astrocytes from pluripotent stem cells. Nat. Biotechnol. 37, 267–275 (2019).

70. Borghese, L. et al. Inhibition of notch signaling in human embryonic stem cell-derived neural stem cells delays G1/S phase transition and accelerates neuronal differentiation in vitro and in vivo. Stem Cells (2010) doi:10.1002/stem.408.

71. Du, Z. W. et al. Generation and expansion of highly pure motor neuron progenitors from human pluripotent stem cells. Nat. Commun. 6, (2015).

72. Cederquist, G. Y. et al. Specification of positional identity in forebrain organoids. Nat. Biotechnol. 37, 436–444 (2019).

73. Baggiolini, A. et al. Developmental chromatin programs determine oncogenic competence in melanoma. bioRxiv 2020.05.09.081554 (2020) doi:10.1101/2020.05.09.081554.

74. Zeng, H. et al. An Isogenic Human ESC Platform for Functional Evaluation of Genome-wide-Association-Study-Identified Diabetes Genes and Drug Discovery. Cell Stem Cell 19, 326–340 (2016).

75. Dzyubenko, E., Rozenberg, A., Hermann, D. M. & Faissner, A. Colocalization of synapse marker proteins evaluated by STED-microscopy reveals patterns of neuronal synapse distribution in vitro. J. Neurosci. Methods 273, 149–159 (2016).

76. Berthold, M. R. et al. KNIME: The konstanz information miner. in 4th International Industrial Simulation Conference 2006, ISC 2006 (2006). doi:10.1145/1656274.1656280.

77. Amin, H. et al. Electrical responses and spontaneous activity of human iPS-derived neuronal networks characterized for 3-month culture with 4096-electrode arrays. Front. Neurosci. 10, (2016).

78. Afgan, E. et al. The Galaxy platform for accessible, reproducible and collaborative biomedical analyses: 2018 update. Nucleic Acids Res. 46, W537–W544 (2018).

79. Patro, R., Duggal, G., Love, M. I., Irizarry, R. A. & Kingsford, C. Salmon provides fast and bias-aware quantification of transcript expression. Nat. Methods 14, 417–419 (2017).

80. Love, M. I., Anders, S. & Huber, W. Differential analysis of count data - the DESeq2 package. Genome Biology (2014).

81. Young, M. D., Wakefield, M. J., Smyth, G. K. & Oshlack, A. Gene ontology analysis for RNA-seq: accounting for selection bias. Genome Biol. 11, (2010).

82. Subramanian, A. et al. Gene set enrichment analysis: A knowledge-based approach for interpreting genome-wide expression profiles. Proc. Natl. Acad. Sci. U. S. A. 102, 15545–15550 (2005).

83. Meers, M. P., Bryson, T. D., Henikoff, J. G. & Henikoff, S. Improved CUT&RUN chromatin profiling tools. Elife 8, (2019).

84. Langmead, B. & Salzberg, S. L. Fast gapped-read alignment with Bowtie 2. Nat. Methods 9, 357–359 (2012).

85. Feng, J., Liu, T., Qin, B., Zhang, Y. & Liu, X. S. Identifying ChIP-seq enrichment using MACS. Nat. Protoc. 7, 1728–1740 (2012).

86. Yu, G., Wang, L. G. & He, Q. Y. ChIP seeker: An R/Bioconductor package for ChIP peak annotation, comparison and visualization. Bioinformatics 31, 2382–2383 (2015).

87. Ramírez, F. et al. deepTools2: a next generation web server for deep-sequencing data analysis. Nucleic Acids Res. 44, W160–W165 (2016).

